# CasTuner: a degron and CRISPR/Cas-based toolkit for analog tuning of endogenous gene expression

**DOI:** 10.1101/2022.10.05.511019

**Authors:** Gemma Noviello, Rutger A. F. Gjaltema, Edda G. Schulz

## Abstract

Certain cellular processes are dose-dependent, requiring a specific quantity of gene products or a defined stoichiometry between them. This is exemplified by haploinsufficiency or by the need for dosage compensation for X-linked genes between the sexes in many species. Understanding dosage-sensitive processes requires the ability to perturb endogenous gene products in a quantitative manner. Here we present CasTuner, a CRISPR-based toolkit that allows analog tuning of endogenous gene expression. In the CasTuner system, activity of Cas-derived repressors is controlled through a FKBP12^F36V^ degron domain and can thereby be quantitatively tuned by titrating the small molecule degrader dTAG-13. The toolkit can be applied at the transcriptional level, using the histone deacetylase hHDAC4 fused to dCas9, or at the post-transcriptional level, using the RNA-targeting CasRx. To optimise efficiency, inducibility and homogeneity of repression we target a fluorescently tagged endogenous gene, *Esrrb*, in mouse embryonic stem cells. Through flow cytometry, we show that CasTuner allows analog tuning of the target gene in a homogeneous manner across cells, as opposed to the widely used KRAB repressor domain, which exhibits a digital mode of action. We quantify repression and derepression dynamics for CasTuner and use it to measure dose-response curves between the pluripotency factor NANOG and several of its target genes, providing evidence for target-specific dose dependencies. CasTuner thus provides an easy-to-implement tool to perturb gene expression in an inducible, tunable and reversible manner and will be useful to study dose-responsive processes within their physiological context.

## Introduction

Biological processes are often dose-dependent, meaning that they rely not only on the presence or absence, but on defined quantities of specific RNAs or proteins. Such dose sensitivity can arise from the need to maintain the right stoichiometry within a protein complex ^1^. It might also evolve to restrict a process to a certain cell type or spatial position within an embryo (e.g. by sensing a morphogen gradient)^2^. As a consequence, a subset of genes exhibit haploinsufficiency ^1,3^ and a dedicated process has evolved in many species to ensure dosage compensation for X-linked genes between the sexes ^4^.

Notably, the process responsible for X-dosage compensation in mammals, X-chromosome inactivation, is itself controlled in a dosage-sensitive manner. It is restricted to female cells by sensing the two-fold higher dose for X-linked genes in females compared to males ^5,6^. Another example for a gene-dosage sensitive process is the differentiation of pluripotent stem cells into different lineages. Here, relatively small variations in the amount of the pluripotency factor OCT4 (POU5F1) can determine whether mouse embryonic stem cells (mESCs) remain in the pluripotent state or differentiate into trophectoderm or meso-endoderm lineages ^7,8^. Similarly, the precise quantity of the pluripotency factor NANOG is critical for the control of näive and primed pluripotent states both *in vitro* and *in vivo* ^9,10^. Understanding the principles underlying dose-dependent regulation of biological processes is, thus, of critical importance. It is however technically challenging, since it requires the ability to quantitatively modulate protein abundance.

The first systems developed that could potentially allow quantitative control of gene expression were inducible promoters such as the TetON/OFF system controlling overexpression of a gene from cDNA ^11^. However, these systems typically overexpress genes beyond their physiological levels and often show uninduced or “leaky” expression ^12,13^. Moreover, intermediate expression can be difficult to achieve at the single-cell level ^14^. Although more complex circuits have been designed to improve quantitative control of gene expression (tunability) ^15^, recent technological developments, such as conditional destabilising domains (degrons) and Cas9-based approaches seem to be more promising for tuning protein abundance ^16–18^. Since they allow control of endogenous genes, they can operate at physiological expression levels.

CRISPR/Cas9-based epigenome editing relies on a catalytically dead Cas9 (dCas9) fused to an effector domain, which is then targeted to a gene promoter through a single guide RNA (sgRNA) to either repress (CRISPRi) ^19^ or activate (CRISPRa) a target gene ^20^. A more recent alternative are RNA-targeting CRISPR systems, such as CasRx ^21^. Different approaches have been applied to make CRISPRa/i inducible, typically relying on conditional stabilisation or on inducible dimerization between dCas9 and the effector domain ^22^. A disadvantage of the latter approach is a partial gene repression in the absence of effector domain recruitment, because dCas9 itself still occupies the target gene promoter ^19^. Therefore, conditional stabilisation is the preferred option for CRISPRi, where a conditional degron domain is fused to a protein of interest to alter its stability in response to stimuli such as binding of ligands ^18^. Although a series of studies have tested different designs of degron-controlled dCas9 systems, they have mostly been applied to CRISPRa and have not investigated tunability at the single cell level ^17,23–25^. Moreover, the most widely used repressor domain, KRAB, has been shown to operate in a switch-like manner ^26^, which would make this classical CRISPRi system unsuitable to generate homogeneous intermediate expression levels. Since in particular CRISPRi would allow modulating expression levels in a physiological meaningful range, we sought to develop a CRISPR/Cas-based system that is tunable at the single-cell level by varying the concentration of a ligand.

We tested a panel of degron and repressor domains and identified two designs that supported potent and tunable repression of a fluorescently-tagged endogenous gene in mESCs. In this system, which we named CasTuner, a FKBP12^F36V^ degron domain ^27^ controls dCas9 fused to a human histone deacetylase 4 (hHDAC4) repressor domain ^28^ or the RNA-targeting CasRx protein ^21^. For both systems we quantify the dynamic properties of gene repression and derepression. Our data show that, by titrating the quantity of repressor using different concentrations of ligand, CasTuner can quantitatively perturb a gene of interest within physiologically relevant ranges homogeneously across cells. We have thus designed a toolkit that allows rapid, inducible, tunable and reversible gene repression at the transcriptional or post-transcriptional level. We show the applicability of CasTuner in studying the dose-dependent action of NANOG in controlling transcription of its targets.

## Results

### The AID and FKBP12^F36V^ degron domains allow potent control of dCas9 abundance

We sought to create a tool for tuning endogenous gene expression that can be easily applied to any target gene. We reasoned that, by fusing a CRISPR-based artificial repressor with a conditional degron domain, we can titrate its quantity and thereby titrate the expression of a target gene. In the first steps, we aimed at identifying suitable degron and repressor domains.

An ideal degron domain would support a wide dynamic range of repressor levels and its complete removal, when not needed. To compare different degrons, we expressed dCas9 fused to a red fluorescent protein (tRFP) tagged with different degron domains, followed by a P2A site and a blue fluorescent protein (tBFP). In these constructs, dCas9-tRFP and tBFP are transcribed together as a bicistronic unit, but are then translated into two separate proteins (Fig. 1a, top). The effect of the degron domain can be quantified as the ratio between tRFP, as a proxy for dCas9 levels, and tBFP, which should remain constant. Using this platform, we evaluated the performance of 6 different degron domains, each fused N-terminally to dCas9: AID ^29^, mAID ^30^, SMASh ^31^, FKBP12^F36V 27^, ecDHFR ^32^ and ER50 ^33^. As a control we used the same dCas9 construct without any degron (“no-degron control”) (Fig. 1a). All constructs were stably integrated in mESCs through PiggyBac (PB) transposition and a cell population with homogeneous tBFP expression was generated by fluorescence-activated cell sorting (FACS) (Fig. 1a, bottom). For the AID-dCas9 and mAID-dCas9 constructs, OsTIR1, an accessory protein required for Auxin-induced proteasomal degradation is expressed from the same construct within the resistance cassette (Fig. S1a).

**Fig. 1:**
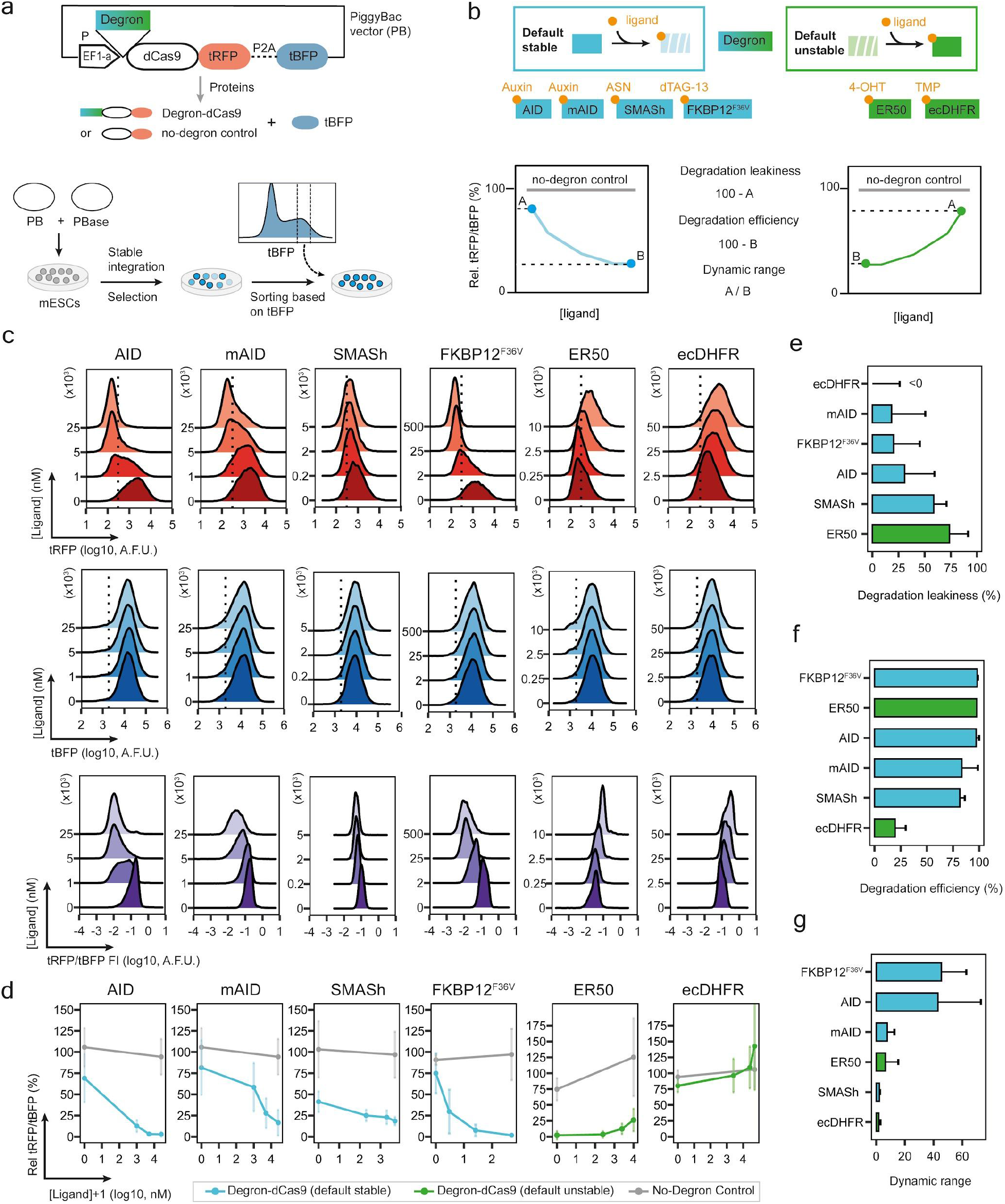
Comparison of degron domains to control dCas9 stability. **a**, Schematic of PiggyBac (PB) plasmids used to assess degron functionality. dCas9 is fused to tagRFP-T (tRFP), followed by a P2A peptide and tagBFP (tBFP) and expressed under the control of EF1-alpha promoter. A degron domain is fused N-terminally to dCas9, while in the “no-degron control” dCas9 is not degron-tagged. The P2A peptide allows cleavage of (degron-)dCas9-tRFP and tBFP into two separate proteins. The PB plasmid is cotransfected with a hyperactive PB transposase (PBase) into mESCs (Tx1072 line). Cells that have genomically integrated the PB plasmid in (possibly multiple) genomic locations are selected with blasticidin and then sorted by FACS based on their tBFP level with identical gates for all constructs, yielding an homogenous population of cells. **b**, Degron-dCas9 titration analysis. Top: overview over the tested degrons, their ligands (orange) and whether they are destabilised (default stable, blue) or stabilised (default unstable, green) by ligand addition. Bottom: based on the tRFP/tBFP ratios measured without ligand, at maximal ligand concentration and for the no-degron control, three parameters are estimated to characterise the ability to control dCas9 for each degron. The tRFP/tBFP ratio was normalised to the tRFP/tBFP ratio in the no-degron control, giving the “relative tRFP/tBFP ratio” (Rel. tRFP/tBFP). We then calculate the degradation leakiness (as measure of the minimal destabilisation conferred by the degron), the degradation efficiency (maximal destabilisation) and the fold change associated as indicated (see Materials and Methods for details). **c**, Degron-dCas9 titration. Density plots for tRFP (top), tBFP (middle) and tRFP/tBFP (bottom) fluorescence intensity (FI) for one biological replicate of each degron-dCas9 line, at increasing ligand concentrations, measured by flow cytometry after 24h of treatment. The tBFP levels are expected to remain constant, while tRFP indicates dCas9 levels. The tRFP/tBFP FI is the ratio of tRFP and tBFP in each cell. Dotted lines show the 99th percentile of non-fluorescent control cells. A.F.U.= Arbitrary Fluorescence Units. **d**, Quantification of degron-dCas9 titration: relative tRFP/tBFP, calculated as Median Fluorescence Intensity (MFI) after subtraction of the background fluorescence and expressed as percentage of the no-degron control at different ligand concentrations. Dots represent the mean of 3 biological replicates and are connected by lines; vertical lines indicate the s.d. **e**-**g**, Bar plots of mean degradation leakiness (**e**), degradation efficiency (**f**) and dynamic range (**g**) calculated for the different degrons as depicted in (**b**). Degron domains from top to bottom are ranked from best to worst for each property. Error bars indicate the s.d. of 3 biological replicates.

We divided the degrons into two groups based on the change in stability of the fusion protein upon ligand addition (Fig. 1b, top): degrons that induce degradation (“default stable”) and degrons that induce stabilisation (“default unstable”). We treated each degron-dCas9 cell line with 4 concentrations of degron-specific ligand for 24 h and measured tRFP and tBFP levels by flow cytometry (Fig. 1b-d and S1b-g). To account for non-specific ligand effects, we also tested the highest concentration of each ligand and the mock treatment on the no-degron control line. For each degron we analysed three different properties (Fig. 1b, bottom): the degradation leakiness, as a measure of destabilisation in the absence of induced degradation (Fig. 1e), the degradation efficiency, which quantifies the destabilisation upon maximal degradation (Fig. 1f), and the dynamic range of the system, describing the fold change between the stabilised and destabilised conditions (Fig. 1g).

Only two out of the 6 tested degron domains were able to efficiently control dCas9 levels, namely the Auxin-controlled AID system and the dTAG-controlled FKBP12^F36V^ degron. Both exhibited a wide dynamic range (43-45 fold change, Fig. 1g), very high degradation efficiency (97-98%, Fig. 1f) and intermediate degradation leakiness (31% and 20%, Fig. 1e). Since FKBP12^F36V^ showed a slightly higher fold change and a lower degradation leakiness than AID, it was employed in all subsequent experiments for post-translational control of Cas-repressor systems.

### The FKBP12^F36V^ degron domain can reliably control Cas-mediated repression of an endogenous gene

Previous work had suggested that the widely used KRAB repressor domain acts in a switch-like manner ^26^ and might therefore not be suitable for quantitative control of gene expression. Thus, we included two additional repression systems in our analyses and compared efficiency, dynamics and homogeneity of repression. We chose dCas9 fused to a histone deacetylase (hHDAC4), which has been shown previously to enable potent repression ^26^, and the RNA-targeting CasRx ^21^. While the KRAB domain induces histone 3 lysine 9 trimethylation (H3K9me3), hHDAC4 catalyses histone deacetylation ^28,34^ and CasRx leads to RNA degradation ^21^ (Fig. 2a). For the dCas9-repressor systems we tested both N- and C-terminal fusion of the repressor domain. All constructs were tagged with the FKBP12^F36V^ degron at their N-terminus and with tBFP at their C-terminus, which allowed monitoring of repressor levels by flow cytometry. For the KRAB-repressor, we created an additional construct, where dCas9 and KRAB are not directly fused, but tethered by the ABA-inducible PYL1/ABI dimerization system (here referred to as KRAB-Split-dCas9) ^35^, with which we have achieved potent repression in the past ^36^.

**Fig. 2:**
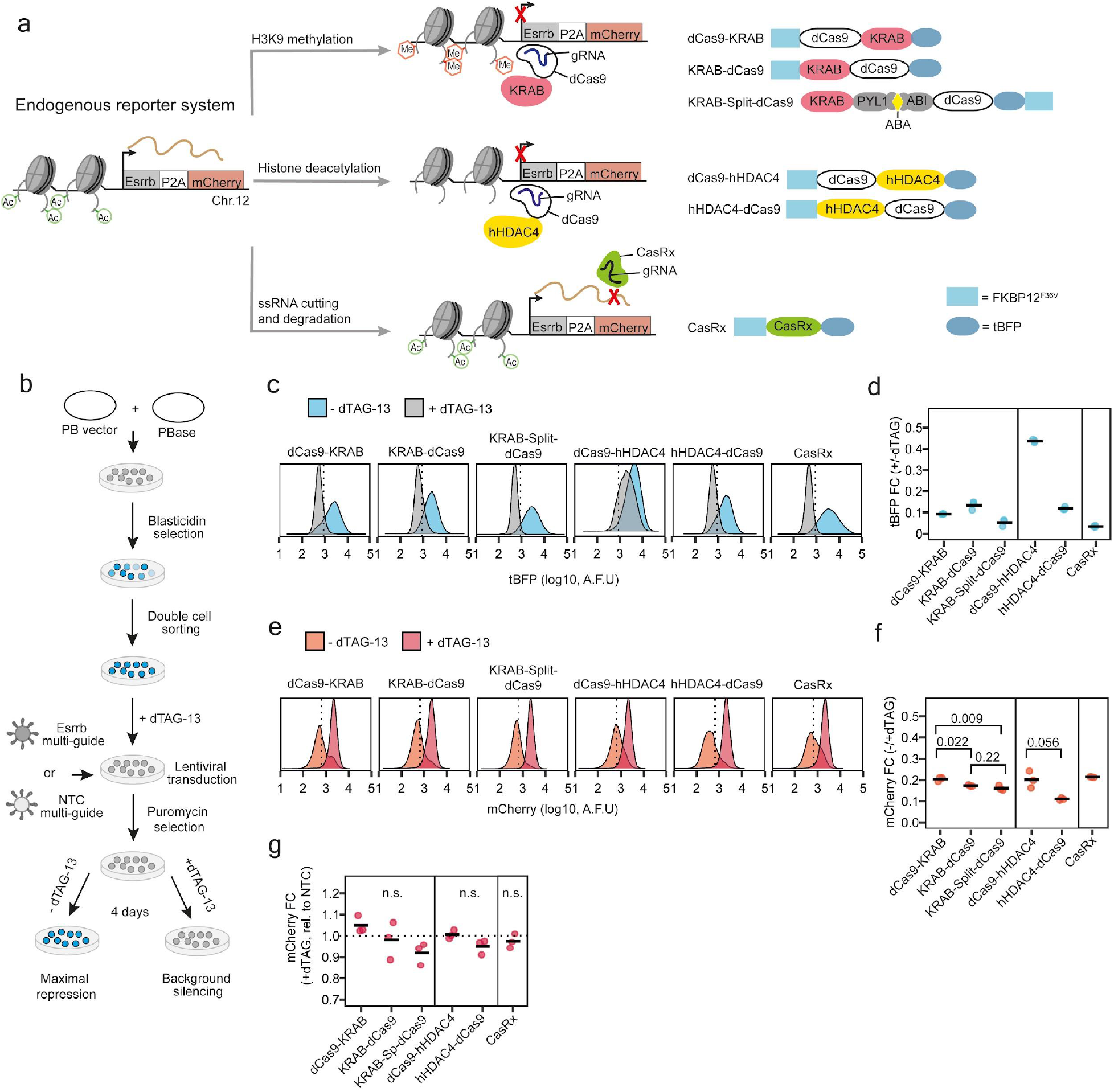
Testing inducibility and efficiency of repression of degron-Cas-repressors in an endogenous reporter system. **a**, Schematic of degron-Cas-repressors and the endogenous reporter system used for their comparison. The *Esrrb* gene is C-terminally fused with P2A-mCherry at its endogenous locus in the 1.8XX mESC line. The *Esrrb* promoter is active in mESCs (green circles= acetyl groups on histone tails) and *Esrrb-P2A-mCherry* sequence is transcribed as a bicistronic unit (brown curved line= mRNA) to then be translated as two separate proteins (ESRRB and mCherry, not shown). Three different repression mechanisms are compared with respect to their ability to repress the reporter gene as indicated. For both KRAB- and hHDAC4-mediated repression, the repressor is tethered to the *Esrrb* promoter region via dCas9. KRAB-mediated repression induces H3K9 methylation (red hexagons) and histone deacetylation, while hHDAC4 induces direct histone deacetylation. Both mechanisms repress transcription. CasRx binds *Esrrb* mRNA via complementary guide RNAs and cuts the transcript causing its degradation. The different designs used for the degron-Cas-repressors are shown on the right. **b**, Experimental strategy for testing inducibility and efficiency of repression of degron-Cas-repressor designs used in (c-g). mESCs cotransfected with PB and PBase plasmids and selected with blasticidin are FACS sorted for two consecutive rounds to ensure uniform expression levels. Cells are then treated with 500nM dTAG-13 to degrade degron-Cas-repressors prior to transduction with lentiviral particles encoding for guide RNAs targeting *Esrrb* or non-targeting control guides (NTC), which are stably integrated. Cells are then either kept in medium with dTAG-13 (+dTAG-13, 500nM) or without dTAG-13 (-dTAG-13, DMSO mock treated) and analysed after 4 days by flow cytometry, to assess background silencing and efficiency of repression, respectively. In parallel to dTAG-13 removal, the KRAB-Split-dCas9 cell line is also treated for 4 days with 100μM ABA to induce tethering of KRAB to dCas9. **c**, Density plots of tBFP levels in cells containing Esrrb-targeting guides, in -dTAG-13 (blue) or +dTAG-13 (grey) conditions, measured by flow cytometry. One biological replicate is shown. The dotted line indicates the 99th percentile of a non-fluorescent control cell line. A.F.U.= arbitrary fluorescence units. **d**, tBFP MFI fold change in +dTAG-13 compared to -dTAG-13 condition in cells with Esrrb-targeting guides. **e**, Same as in (**c**) but for mCherry. **f**, mCherry expression fold change of cells with Esrrb-targeting guides in -dTAG-13 compared to +dTAG-13 condition. The KRAB-dCas9 and KRAB-Split-dCas9 lines showed a small but significantly stronger knock-down of ESRRB compared to dCas9-KRAB and a similar tendency is observed for hHDAC4-dCas9 compared to dCas9-hHDAC4. The values of unpaired t-tests are reported. **g**, mCherry expression fold-change of cells with Esrrb-targeting guides compared to NTC guides in +dTAG-13 conditions. No difference was observed between targeting and non-targeting guides (p-value non significant, unpaired t-test). In (**d**,**f**,**g**) 3 biological replicates are shown as dots with a horizontal bar showing their mean.

To compare their ability to tune endogenous gene expression, we targeted the repressor systems to the *Esrrb* gene in a mESC line, where the gene is homozygously tagged with P2A-mCherry (Fig. 2a) ^36^. We then used flow cytometry to quantify repression at the single-cell level. Previous work had shown that mESCs lacking *Esrrb* retain self-renewal and appear morphologically normal ^37^, making it a suitable reporter system to test different repression mechanisms on endogenous gene expression.

For each repressor construct, we created stable cell lines through PB transposition followed by two consecutive rounds of cell sorting based on tBFP levels, in order to obtain cells with homogeneous repressor abundance (Fig. 2b). We then transduced cells with a sgRNA vector, co-expressing three different guide RNAs targeting the *Esrrb* promoter or expressing non-targeting control (NTC) guides. Transduction was performed in the absence of the targetable repressor protein (high dTAG-13). To induce repression, the dTAG-13 degrader was withdrawn for 4 days (Fig. 2b). We then measured ESRRB expression (mCherry) and the quantity of repressor (tBFP) by flow cytometry (Fig. 2c-f and S2).

All degron-Cas-repressors were efficiently degraded with the exception of the C-terminal hHDAC4 construct (dCas9-hHDAC4), where tBFP levels were only reduced by ∼56% (Fig. 2c-d). When comparing mCherry levels in the presence and absence of dTAG-13, the N-terminal fusion of the repressor domain resulted in strong depletion for both KRAB and hHDAC4 fusions (∼83-89%), which was somewhat weaker for the C-terminal fusion constructs (∼80%) (Fig. 2e-f). CasRx also induced a clear knock-down with 79% mCherry reduction. The strength of repression did not strictly depend on the repressor level, since FKBP12^F36V^-dCas9-hHDAC4 was expressed at higher levels compared to FKBP12^F36V^-hHDAC4-dCas9, but induced less repression (Fig. S2f). To assess whether residual degron-Cas-repressor protein in the presence of dTAG-13 would result in unwanted repression, we compared mCherry levels between cells transduced with a targeting and a non-targeting sgRNA construct. We could not detect any difference between the two lines, which showed that repression was effectively prevented by repressor degradation and no repression leakiness was observed (Fig. 2g).

In summary, we have generated a set of degron-controlled Cas-repressor constructs that can efficiently repress a target in a strictly inducible manner. We found an optimised design, where the repressor domain is fused to the N-terminus of dCas9, which confers increased repressor activity and will be characterised further in the next sections.

### HDAC4- and CasRx-mediated repression allows analog tuning of gene activity

Having optimised several degron-Cas-repressor systems, we next aimed at comparing their ability to tune endogenous gene expression at the single-cell level. We refer to repression “tunability” as the ability to partially repress a target gene, resulting in stable intermediate expression levels. Our goal is to homogeneously titrate protein abundance by controlling the quantity of repressor. Depending on the repression mechanisms, intermediate repressor levels could in principle lead to two alternative outcomes: a gradual and homogenous change in target gene expression or a bimodal pattern with a positive and a negative population of cells. Hence, the modality of repression can be defined as “analog” in the first scenario and “digital” in the second (Fig. 3a).

**Fig. 3:**
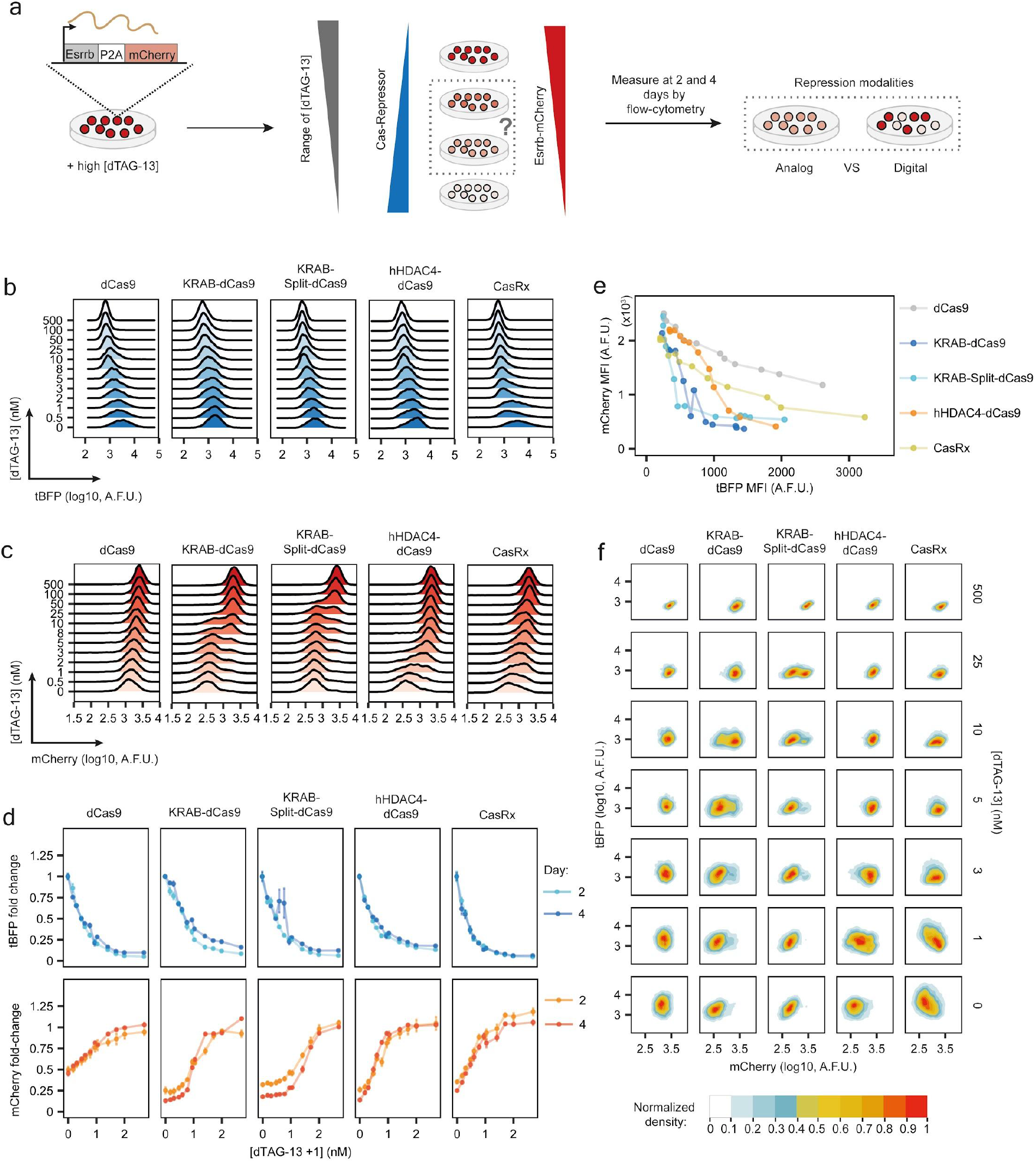
Assessing tunability of degron-Cas-repressor systems. **a**, Experimental design: degron-Cas-repressor cell lines expressing ESRRB-mCherry are transferred from medium containing high concentration of dTAG-13 (repressor degraded) to media with a range of dTAG-13 concentrations. The degron-Cas-repressor levels are expected to decrease with increasing dTAG-13 concentrations, resulting in a rise in target gene expression (mCherry). Quantification of ESRRB-mCherry and degron-Cas-repressor (tBFP) levels after 2 and 4 days by flow cytometry will then allow the distinction between analog (homogenous intermediate levels) and digital (a mixture of positive and negative cells) repression. **b, c**, Density plots of tBFP (**b**) and mCherry (**c**) expression levels measured by flow cytometry at increasing dTAG-13 concentrations (y-axis, in nM), for degron-Cas-repressors as indicated on top, expressing Esrrb-targeting guides after 4 days of treatment. One biological replicate is shown. dCas9 and KRAB-Split-dCas9 are the same construct (KRAB-Split-dCas9) with 100µM ABA being added to the KRAB-Split-dCas9 4 days before the measurement. **d**, tBFP (top) and mCherry (bottom) fold change of background-subtracted MFI in cells expressing Esrrb-targeting guides, at different dTAG-13 concentrations (y-axis, log10 +1 scaled, in nM) after 2 or 4 days of treatment, normalised to mean MFI in cells expressing NTC guides treated with the same dTAG-13 concentrations. The mean of three biological replicates (dots) ± standard deviation (vertical bars) is shown. Lines connect the means. **e**, mCherry MFI at different doses of repressor (tBFP MFI). The mean of three biological replicates is shown. **f**, Normalised 2D density plots showing tBFP (y-axis) and mCherry (x-axis) levels in populations of cells treated with different dTAG-13 concentrations (indicated on the right, in nM). A.F.U. = Arbitrary Fluorescence Units

We used our endogenous Esrrb-mCherry reporter system and titrated each degron-Cas-repressor through treatment with varying dTAG–13 concentrations. We characterised CasRx and the dCas9 constructs with N-terminal repressor domains, but also the effect of dCas9 alone (KRAB-Split-dCas9 without ABA treatment), which likely acts through steric hindrance of transcription.

By using a range of 12 dTAG-13 concentrations, we were able to homogeneously titrate protein abundance (tBFP) of all 5 repressors (Fig. 3b, S3). When analysing target gene expression (Fig. 3c-d), we observed that KRAB-mediated repression was effective at lower repressor levels than hHDAC4 and CasRx (Fig. 3e). The KRAB domain thus appears to be the most potent repressor of the three. When analysing tunability at intermediate repressor levels we observed two different repression patterns: CasRx, dCas9 alone and dCas9-HDAC4 exhibited gradual tuning of mCherry levels, while KRAB-mediated repression gave rise to a bimodal distribution (Fig. 3c). Comparison of mCherry-high and -low cells at intermediate KRAB repression strength revealed that both populations expressed similar amounts of tBFP (Fig. 3f). The observed bimodal repression pattern is thus unlikely to arise from heterogeneity in repressor levels, but appears to be an inherent property of the repression mechanism. Clearly, repression tunability can be verified only by single-cell measurements, since at the cell population level, bimodal or homogeneous intermediate distributions are indistinguishable (Fig. 3c vs 3d). Among the tested systems, only hHDAC4-dCas9 and CasRx are able to tune gene expression at the single-cell level. Therefore, we named these two systems collectively the CasTuner toolkit.

### Speed and reversibility of repression in degron-Cas-repressor systems

Having identified several degron-Cas-repressor designs that can quantitatively tune gene expression, we set out to further characterise their dynamics and reversibility. In addition to the tunable CasRx and hHDAC4-dCas9 systems of the CasTuner toolkit we again included the widely used KRAB systems and dCas9 alone for comparison. The dynamics of repression upon ligand withdrawal and derepression upon ligand addition depend on the repressor dynamics, but also on the speed with which the repressed state is established and erased, respectively. For CasRx we expect establishment and erasure of repression to be immediate, since here the repressor itself will cleave the mRNA. For HDAC4 and KRAB-dependent systems these processes might be slower, since they affect gene expression in a less direct manner through modifying the chromatin state at the gene promoter.

To assess the repression and derepression dynamics, we induced repressor upregulation by dTAG-13 withdrawal and repressor degradation by dTAG-13 addition, respectively. In each case we then monitored repressor (tBFP) and target gene (mCherry) levels at different time points over 6 days (Fig. 4, Fig. S4a-d). To disentangle the different steps that control the system’s dynamics, we parameterized ordinary differential equation (ODE) models of the system based on the collected data set (Fig. S4e-g). For each system we estimated two different parameters: the time required to reach half of the final repressor level (t_1/2_) and the delay between repressor up- or downregulation and effects on target gene expression (Δt). We assumed that the repressors were produced at a constant rate and degraded in a dTAG-13-dependent manner. For mCherry the production rate was assumed to be a function of the repressor level and the associated parameters were estimated from the dTAG-13 titration experiments (Fig. S3e). The mCherry degradation rate was assumed to be identical in all cell lines and was estimated from the derepression time course for CasRx under the assumption that release of repression was immediate in this system (Fig. S4e, see Methods section for details).

**Fig. 4:**
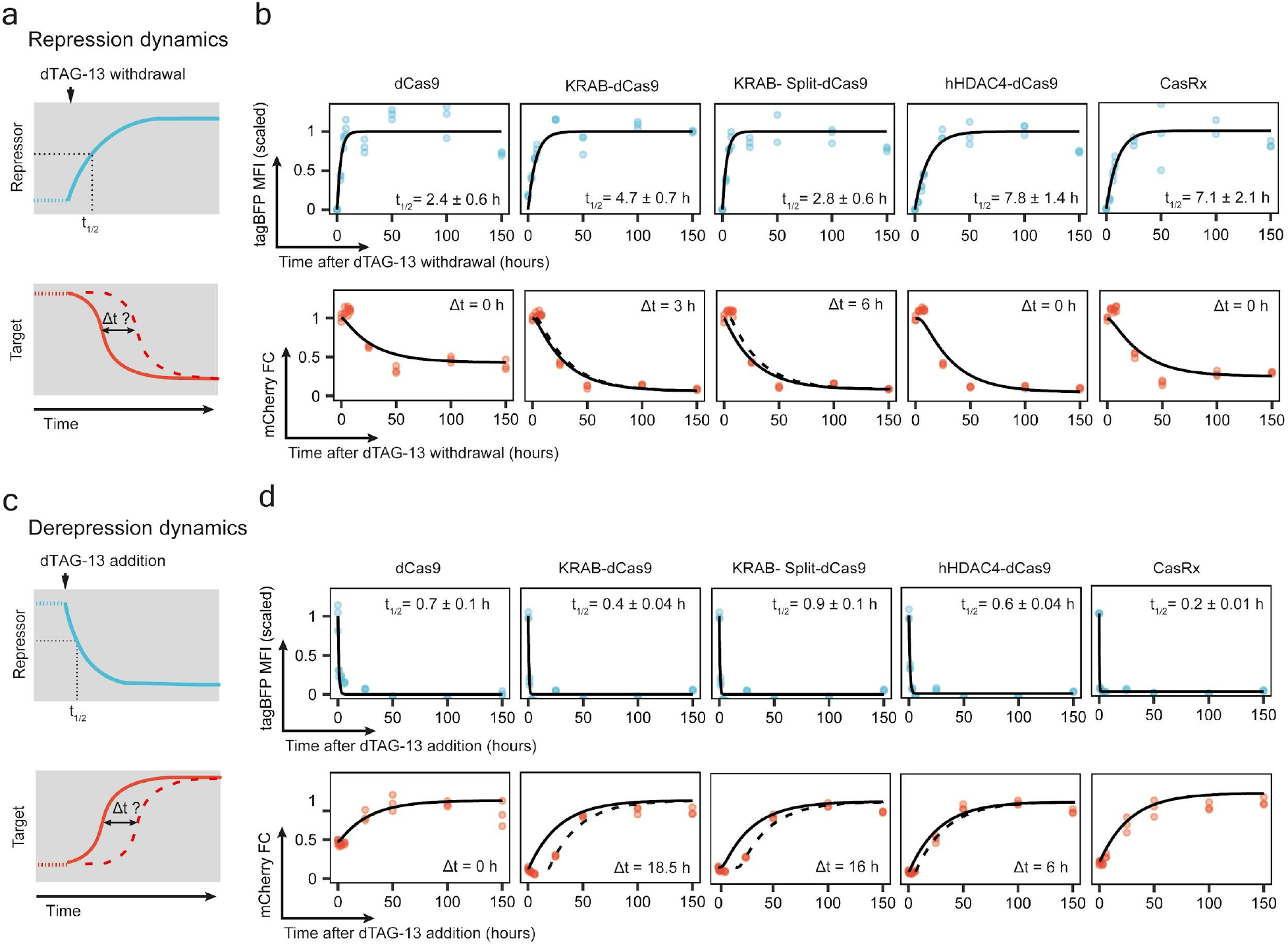
Repression and derepression dynamics of degron-Cas-repressor systems. **a**, Schematics of experiment to study the dynamics of repression for the different degron-Cas-repressors. dTAG-13 is withdrawn at time 0 and then repressor (tBFP) and target (ESRRB-mCherry) levels are measured by flow cytometry over time. Upon ligand withdrawal, the degron-Cas-repressor level increases to then reach a steady state. We can estimate the time required to reach half of the maximal repressor level (t_1/2_) by fitting an ordinary differential equation (ODE) to the experimental data. To assess whether target repression is immediate upon repressor upregulation (Δt=0, solid line) or occurs with a delay (Δt>0, dashed line), the mCherry data was fitted with an ODE model, where target gene expression varies as a function of time and of repressor concentration with or without assuming a delay between repressor upregulation and target gene repression (Δt). See Fig. S4e and the methods section for details on the modelling approach. **b**, Repressor (tBFP) MFI, scaled between 0 and 1 and target gene (mCherry) fold change (FC) for the different degron-Cas-repressor systems, for 3 biological replicates (dots), at different time points in the time-course experiment to study the repression dynamic schematized in (a). The black lines in each plot represent the best fit of the ODE model. For tBFP, the t_1/2_ ± its standard error is indicated for each degron-repressor system. For mCherry, the continuous line shows the result of the ODE model with Δt = 0 h and the dashed line the ODE model with the Δt that minimises the Mean Absolute Error of the model compared to the experimental measurements. **c**, To study the dynamic of derepression associated with each system, after 4 days in the absence of dTAG-13 (in the presence of repression), dTAG-13 is added back to the medium. The same type of model shown in (**a**) is used and we estimate the delay between repressor degradation and target gene derepression. **d**, Same as in (**b**) but for the derepression dynamics experiment schematized in (**c**). Both KRAB-based repression systems show a substantial delay in target gene derepression.

The time required to reach half of the maximal repressor level upon ligand withdrawal (t_1/2_) ranged from ∼3h for dCas9 and KRAB-Split-dCas9 to ∼7-8 h for CasRx and hHDAC4 (Fig. 4b top). These differences are likely due to variation in protein turnover rates in the absence of ligand, because those determine the dynamics of upregulation. Once the repressor was upregulated, repression was immediate for dCas9, hHDAC4-dCas9 and CasRx (Δt=0h), while a small delay was detected for the two KRAB-mediated systems (Δt=3-6h, Fig. 4b bottom). Hence, our analyses show that the speed of target depletion is mainly determined by the repressor dynamics (and target stability) and that full repression is established within two days for all systems.

When analysing derepression upon dTAG-13 addition after 4 days of culture without the degrader, all constructs were rapidly depleted with a half time of <1h (Fig. 4d, top). Although repression was fully reversible in all cases, we observed clear differences in the dynamics. While derepression was immediate for dCas9 alone, a delay of 16-18h was observed for the KRAB systems, which was substantially smaller (6h) for hHDAC4-dCas9 (Fig. 4d, bottom). In summary, we have estimated the dynamics of repression and derepression in our reporter system for different degron-Cas-repressors. While target repression appears to be immediate, derepression dynamics depend on the mode of repression.

### CasTuner can quantify dose-response curves between NANOG and its target genes

We designed CasTuner in order to finely regulate the quantity of a target gene. This allows us to study how gene dose relates to a phenotype of interest (dose-response). As a proof of concept, we applied CasTuner to titrate the dose of the core pluripotency factor NANOG in mESCs and measured how its quantity affects expression of known target genes (Fig. 5a). Since NANOG safeguards the pluripotent state ^38^, a threshold level of NANOG quantity for induction of differentiation has been proposed to exist ^39^, raising the question of how NANOG dose controls such a process.

**Fig. 5:**
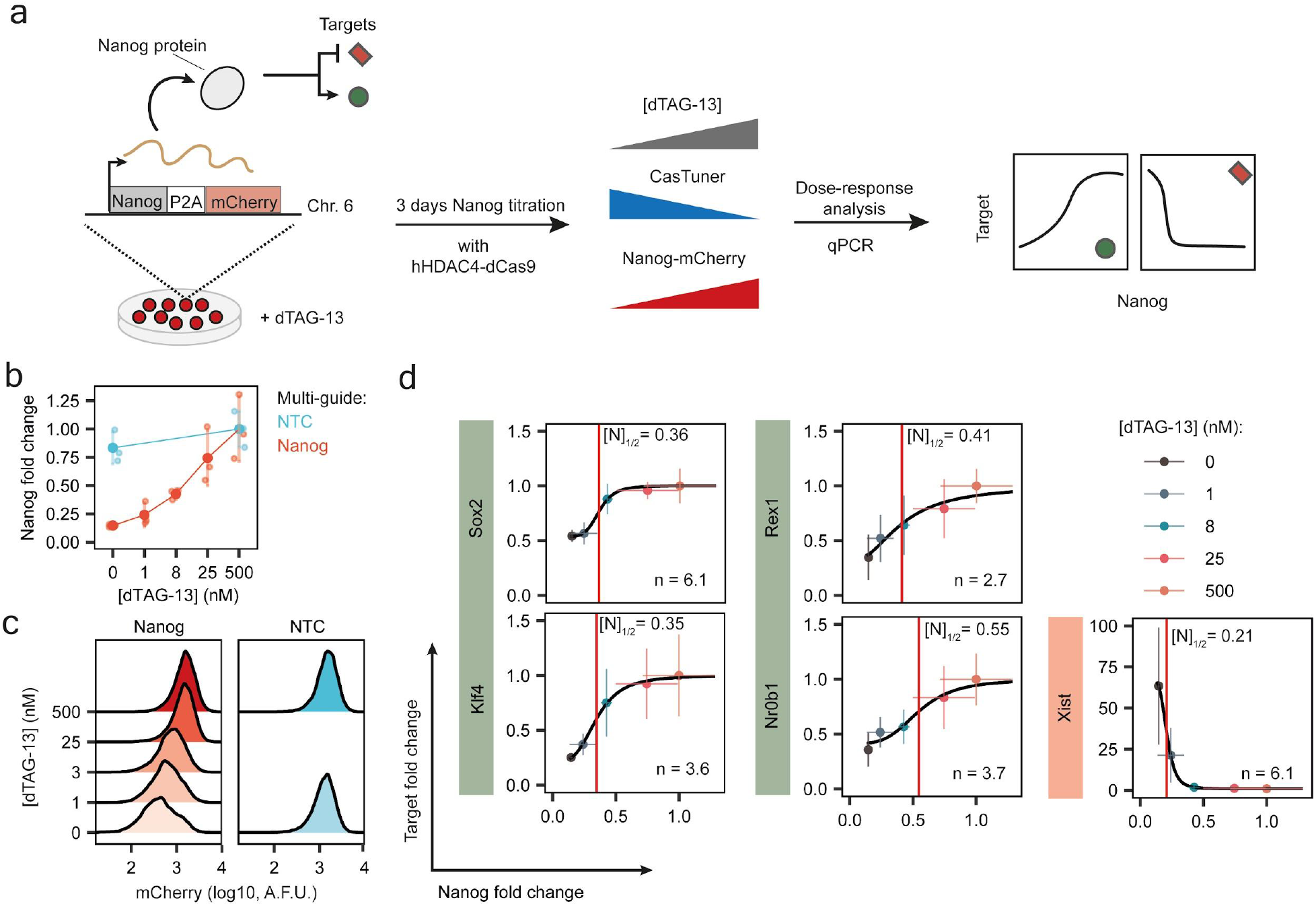
NANOG dose-response curves measured using CasTuner. **a**, Experimental strategy to measure dose-response curves between NANOG and its target genes. A mESC line with *Nanog* fused with P2A-mCherry at its endogenous locus, stably expressing CasTuner (FKBP12^F36V^-hHDAC4-dCas9) is used. After transduction with Nanog-targeting or non-targeting guides in the presence of dTAG-13 (500nM, no repression) the cells were treated with a range of dTAG-13 concentrations in order to titrate NANOG. RNA was harvested after 3 days to quantify *Nanog* and its target genes by qPCR. NANOG levels are expected to positively correlate with targets activated by NANOG and negatively with repressed targets. **b**, *Nanog* fold change at different dTAG-13 concentrations (x-axis, in nM) compared to cells with NTC guides at 500nM dTAG-13, as measured by qPCR, for 3 biological replicates (small dots); vertical lines indicate the standard deviation and big dots the mean. **c**, mCherry levels as a proxy for NANOG protein measured by flow cytometry in cells expressing Nanog-targeting or NTC guides, treated with the indicated dTAG-13 concentrations. Only one replicate was measured and the experiment was performed in the exact same conditions as in (b). AFU=arbitrary fluorescence units. **d**, Dose-response curves for NANOG target genes at varying *Nanog* levels, expressed as fold change of relative expression compared to the 500nM dTAG-13 condition. Shown as the mean of 3 biological replicates (dots). Horizontal and vertical lines indicate the standard deviation for *Nanog* and its target genes, respectively. The black curve represents the most likely Hill curve fitted using a non-linear least square approach (see Materials and Methods). The red vertical line marks the fold change of *Nanog* that leads to half of the maximal expected fold change variation of the target ([N]_½_), according to the fitted model, which is also reported in each plot. The R^2^ for the fits ranges between 0.94 and 0.99.

We employed an mESC line (1.8XX-Nanog-mCherry) where the endogenous *Nanog* gene is homozygously tagged at its C-terminus with P2A-mCherry ^36^. We stably integrated the FKBP12^F36V^-hHDAC4-dCas9 system by PB transposition and generated a cell population with homogenous CasTuner expression through two rounds of cell sorting based on tBFP levels. The cells were transduced in the absence of CasTuner expression (high dTAG-13) with a lentiviral multi-guide plasmid targeting the *Nanog* promoter region or with NTC guides. We initially treated cells with 12 different dTAG-13 concentrations and analysed mCherry levels after 1, 2, 3 and 4 days by flow cytometry (Fig. S5a-d). While repression was clearly visible after 1 day (∼3 fold reduction), the strongest depletion was observed between 2 and 3 days (∼8 fold). Based on these data we chose 5 dTAG-13 concentrations, for which we analysed target gene expression after 3 days by qRT-PCR (Fig. 5). We confirmed that NANOG was indeed titrated both at the mRNA and protein level (Fig. 5b-c). We then quantified the expression of NANOG target genes *Rex1* (Zfp42), *Nrb01, Sox2* and *Klf4*, which have been reported to be activated by NANOG ^40^, and of *Xist*, which is known to be repressed by NANOG ^41^. As expected *Xist* was upregulated upon *Nanog* repression, while the other targets were reduced (Fig. 5d). We observed marked differences in the dose-response curves between the tested targets. To better characterise the dose-response relationships, we fitted a Hill curve for each target gene. In this way, we estimated two parameters: (1) the “target sensitivity” N_1/2_, given by the NANOG reduction required to induce half of the maximally observed target down- or upregulation, and (2) the degree of non-linearity of the response given by the Hill coefficient “n”, where a high value (n>>1) indicates a switch-like response, and a lower value a more gradual mode of regulation (see Materials and Methods). Among the tested genes, *Nr0b1* reacted most sensitively with N_1/2_=0.55, while *Xist* required a much stronger NANOG reduction to be affected (N_1/2_=0.21). Moreover, we found that *Sox2* and *Xist* were controlled in a more switch-like manner with n∼6, while the other targets reacted more gradually (n=2.2-3.8). These results suggest that different NANOG doses can affect target gene expression in different ways. Therefore, a unique NANOG threshold does not seem to exist, pointing to a more nuanced picture of NANOG-mediated pluripotency regulation.

## Discussion

In this study we have developed a novel toolkit, CasTuner, based on Cas proteins fused with a conditional degron domain. CasTuner allows quantitative control of endogenous gene expression either at the transcriptional (with hHDAC4-dCas9) or post-transcriptional level (with CasRx). The system employs the FKBP12^F36V^ degron to precisely control Cas repressor abundance by varying the concentration of the dTAG-13 ligand. We show that intermediate CasTuner levels can homogeneously tune target gene expression in an analog manner, as opposed to the widely used KRAB-mediated CRISPRi system, which exhibits a digital mode of action. Moreover, we show that in mESCs, repression dynamics are determined only by the repressor stability and that repression is readily reversed upon ligand addition, with derepression occurring more rapidly with CasTuner compared to the KRAB system. Since CasTuner can titrate endogenous gene activity in a physiologically relevant range, the toolkit will allow us to gain a better understanding of how cellular functions are quantitatively controlled in mammals.

As a proof-of-concept we employed CasTuner to titrate the core pluripotency factor NANOG in mESCs and measured dose-response curves for a set of NANOG target genes. We show that sensitivity and also non-linearity of the response is variable between targets. An extreme scenario is found for *Xist*, which is repressed by NANOG in mESCs ^41^ and upregulated in female cells upon differentiation. Xist upregulation required ∼80% Nanog depletion, while other targets were already affected by a change of less than 50%. Different scenarios could be envisioned to explain these observations. Nanog might bind its target genes with varying affinity, such that a subset will remain occupied even at lower concentrations. This could be due to differences in sequence composition or spatial arrangements of binding sites. In particular the latter case could lead to stark differences in occupancy, because NANOG has been shown to oligomerize *in vivo* ^42^. Alternatively, NANOG might co-bind with other factors that are also controlled by NANOG at the transcriptional level, which could lead to differential sensitivity depending on which cofactor is employed. For Xist, where NANOG binds together with multiple other factors including SOX2 ^41,43^, repression might for example only be released, once NANOG and SOX2 are depleted, which only occurs at lower NANOG levels. Our results reveal that certain repression mechanisms, such as histone deacetylation and RNA degradation function in an “analog” way in the sense that they can induce stable intermediate expression levels when analysed with single-cell resolution. Others, such as KRAB-mediated systems, work in a “digital” manner, where the quantity of the repressor only defines the percentage of cells that will shut off target gene expression. In the latter case, bimodal distributions are observed for the targeted gene at intermediate repressor levels. Such bimodal patterns are typical for bistable systems, which usually respond in a switch-like manner ^44^. In the context of gene regulation, bistability can arise from positive feedback loops in chromatin regulation ^45^. Here histone-modifying enzymes are recruited through binding the modification they deposit either directly or in an indirect manner through additional proteins. Such feedbacks have been found in particular for repressive modifications, such as H3K27me3, which is deposited by the PRC2 complex ^46^, and H3K9me3, which is catalysed by several enzymes in mammalian cells, including SETDB1, which is recruited by the KRAB repressor domain via a KAP1-mediated interaction ^47^. Such a mechanism is also thought to allow spreading of the repressed state along the DNA to form chromatin domains, and to underlie epigenetic memory ^34,45,48^. Although silencing is readily reversible in mESCs for all repressors we tested, the observed slower derepression for the KRAB systems might be due to bistability, which seems to confer (short-term) epigenetic memory. Notably, derepression dynamics appear to be substantially slower in other more differentiated cell types ^26,49,50^. While KRAB systems induce a repressed chromatin state, hHDAC4 in the CasTuner toolkit functions via removal of active histone modifications, namely acetylation of a multitude of histone residues ^51^. Erasure of the active chromatin state at target gene promoters thus appears to enable analog tuning as opposed to induced heterochromatization.

The CasTuner toolkit also provides a chromatin-independent approach to tune expression levels, where RNA degradation is manipulated through the CasRx system. This approach is similar to direct degron-tagging of a target gene, since both modulate protein stability. Indeed, our data shows that direct tagging with the FKBP12^F36V^ degron also allows analog tuning of the tagged protein (tBFP-tagged Cas repressor in our case). Therefore direct degron-tagging of the target gene would be an alternative approach to homogeneously titrate protein levels, which would allow faster response dynamics compared to CasTuner, where the response time is limited by the repressor half-life. Direct degron-tagging is however associated with some major drawbacks, such as the often-observed basal (uninduced) degradation ^52^, possible impediments in structural folding and interactions and labour-intensive generation of the required cell lines through gene targeting. With CasTuner, titration of a new target gene requires only a single guide-encoding plasmid. The approach is therefore relatively simple and cost-effective, allowing the study of how the quantity of different genes relates to a phenotype, thus preserving the scalability that characterise CRISPR-based technologies.

In the future, CasTuner could also be expanded to an orthogonal system. Since we identified two potent degron domains (AID and FKBP12^F36V^) and two tunable repressor systems (dCas9-hHDAC4 and CasRx), which employ different types of guide RNAs, each repressor could be controlled by a different degron tag. In this way, treatment with variable concentrations of different degrader molecules would allow simultaneous independent tuning of two endogenous genes. An interesting application for CasTuner and also for a CasTuner-based orthogonal system could be the quantitative analysis of phase separation phenomena in cells. Although phase separation has been suggested as regulatory mechanisms for a variety of cellular processes, its occurrence and functional importance *in vivo* is often debated ^53^. A hallmark of phase separation is the existence of a saturation concentration for the involved macro-molecules, above which phase separation occurs ^54^. Testing the existence of a saturation concentration in cells requires the ability to titrate macro-molecules such as proteins or RNAs *in vivo*. Here CasTuner would be a powerful tool to address this technical challenge in the growing phase separation field.

## Materials and Methods

### Cell lines

The female TX1072 mESC line is a F1 hybrid ESC line derived from a cross between the 57BL/6 (B6) and CAST/EiJ (Cast) mouse strains that carries a doxycycline-responsive promoter in front of the *Xist* gene on the B6 chromosome and an rtTA insertion in the *Rosa26* locus ^55^. The 1.8XX Nanog-mCherry and Esrrb-mCherry reporter lines are female mESC lines that carry a homozygous insertion of 7xMS2 repeats in *Xist* exon 7 and a C-terminal P2A-mCherry tag at the *Nanog* or *Esrrb* genes, respectively ^36^.

TX1072 mESC lines with degron-dCas9-tRFP-P2A-tBFP constructs and 1.8XX Nanog-mCherry and Esrrb-mCherry reporter lines expressing degron-Cas-repressor-tBFP constructs were generated through piggyBac transposition, antibiotic selection with blasticidin and FACS based on tBFP fluorescent levels (see below).

### mESCs culture

All mESC lines were grown without feeder cells on gelatin-coated flasks (Millipore, 0.1%). mESCs were passaged every second day at a density of 4 × 10^4^ cells/cm^2^ and medium was changed daily. Cells were grown in serum-containing medium (DMEM (Sigma), 15% ESC-grade FBS (Gibco), 0.1 mM β-mercaptoethanol), supplemented with 1000 U/ml leukaemia inhibitory factor (LIF, Millipore) only for all experiment performed in 1.8XX mESCs or supplemented with LIF and 2i (3 μM Gsk3 inhibitor CT-99021, 1 μM MEK inhibitor PD0325901, Axon), when growing TX1072 cell lines. For the NANOG titration experiment the cells were seeded at a lower density (3 × 10^4^ cells/cm^2^) and not passaged before RNA harvesting, to counteract possible selective effects of *Nanog* knock-down. For experiments with a flow cytometry readout, cell treatment and analysis was usually performed in 96-well plates. Here cells were seeded at a density of 20,000 cells per well and were passaged 1:8 after two days for longer treatments.

### PiggyBac transposition

In order to generate cell lines stably expressing dCas9 and CasRx constructs, expression plasmids were genomically integrated through piggyBac transposition. To this end mESCs were transfected using Lipofectamine™ 3000 Transfection Reagent (Invitrogen) with the donor plasmid and a plasmid encoding for a hyperactive piggyBac transposase (pBROAD3-hyPBase-IRES-zeocin, a kind gift from the Giorgetti lab) in a 5:1 molar ratio using a total of 2.5µg of DNA. A reverse transfection protocol was employed, where 0.4 × 10^6^ cells are seeded together with the lipofection mixture containing the plasmids in a 6-well plate coated with gelatin. The lipofection mixture was prepared according to the manufacturer’s instructions. On the next day, fresh medium was added to the cells. On the second day cells were transferred to a T25 flask with medium containing blasticidin (5ng/µl, Roth), followed by selection for 7 days. During antibiotic selection, the medium was changed daily and cells were passaged when they became ∼80% confluent, to favour an even selection of all cells.

### FACS

For Fluorescence Associated Cell Sorting (FACS), a BD FACSAria Fusion sorter (Beckton Dickinson, IC-Nr.:68198, Serial-Nr.:R658282830001) with a 2B-5YG-3R-2UV-6V lasers configuration was used. The cells were sorted based on their tBFP level. For an example of the strategy employed to sort cells for the testing of different degrons fused to dCas9, see Fig.1A. An example of the gating coordinates used for double sorting of cells with degron-Cas-repressor systems is given in Supplementary File 1. The strategy for sorting was selected in order to obtain a high number of cells, with an as uniform as possible level of expression, while maintaining high expression levels of the construct, clearly distinguishable from the fluorescent background of the cells. Immediately after sorting, the cells were centrifuged, resuspended in the appropriate medium with 1x Penicillin-Streptomycin (Gibco, #15070063) and seeded. Penicillin-Streptomycin was kept for 2 passages.

### Lentiviral transduction

For the generation of cell lines carrying CRISPRi or CasRx multi-guide plasmids, we used lentiviral transduction. DNA constructs were first packaged into lentiviral particles. For this, 1 × 10^6^ Hek293T cells were seeded into one well of a 6-well plate and transfected the next day with the lentiviral packaging vectors: 1.2 μg pLP1, 0.6 μg pLP2, and 0.4 μg VSVG (Thermo Fisher Scientific), together with 2 μg of the desired construct using Lipofectamine 2000 (Thermo Fisher Scientific). Hek293T supernatant containing the viral particles was harvested after 48 h. The viral supernatant was concentrated using Lenti-X™ Concentrator (#631232, Takara) according to the manufacturer’s instructions, resuspended in 200µl of mESC medium supplemented with LIF and frozen at −80°C in 50µl single use aliquots. 0.2 × 10^6^ mESCs were seeded per 12-well and transduced the next day with 50 μl of concentrated viral supernatant and 8 ng/μl polybrene (Sigma). Antibiotic selection was started 2 days after transduction and kept until all cells in the non-transduced control were dead (typically within 1-2 passages).

### Ligands

Auxin (3-Indoleacetic acid, IAA, GoldBio #I-110-25) was dissolved in EtOH to a 400mM stock dilution, according to the manufacturer instructions. Aliquots were kept at −20°C, protected from light. dTAG-13 (Tocris #6605) was dissolved in DMSO to a 5mM stock concentration, according to manufacturer instructions. Aliquots were kept at −20°C. Trimethoprim (TMP, Sigma #T7883) was dissolved in DMSO to a concentration of 200mM and stock aliquots were kept at −20°C. Asunaprevir (ASN, MedChem Express #HY-14434) was dissolved in DMSO to 10mM stock solution and stored at −20°C. (Z)-4-Hydroxytamoxifen (4OHT, Sigma #H7904) was dissolved in EtOH to a stock solution of 10mM and stored at 4°C.

### sgRNAs design

sgRNAs to target the *Esrrb* and *Nanog* promoters were designed using the CRISPR-Cas9 online tool Chopchop (https://chopchop.cbu.uib.no/) ^56^. Because *Esrrb* has multiple isoforms with different transcription start sites, the mESC-specific isoform (ENSMUST00000115313.7) was used in the query. Four of the highest ranking guides were selected, avoiding to pick guides in close proximity between each other (<50 base pairs) and/or containing BsmBI restriction sites. Non-targeting control (NTC) sgRNAs were extracted from previous publications ^57^ and are predicted to target non-functional genomic regions.

For CasRx-mediated repression, the online tool https://cas13design.nygenome.org/ was used to design guides targeting *Esrrb* ^58^. The shortest translated transcript isoform that contains all constitutive exons was used in the query (ENSMUST00000167891.1) and the 3 highest ranking guide sequences were selected. As negative controls we extracted safe-targeting control guide sequences from a previous publication and removed 4 nucleotides at the 5’-end ^59^, to obtain 23 base pairs long sequences. The resulting NTC sequences were confirmed to have low similarity with sequences contained in the NCBI Transcript Reference Sequences database for mus musculus, using Blast https://blast.ncbi.nlm.nih.gov/Blast.cgi. The guide sequences used in this study are provided in Supplementary table 1 (part A for dCas9 and part B for CasRx).

### Molecular cloning

All cloned plasmids described in this study were transformed in NEB^®^ Stable Competent *E. coli* (High Efficiency) (#C3040H). Plasmids isolated from selected colonies were analysed by Sanger sequencing to confirm successful cloning.

#### sgRNAs cloning in multi-guide expression system

Multi-guide plasmids for CRISPRi were cloned as previously described ^36^. Briefly, four different sgRNAs were cloned into the sgRNA expression plasmid SP199 ^36^, with Golden Gate cloning, such that each sgRNA is controlled by a different Pol III promoter (hU6, mU6, 7SK and hH1).

#### CasRx guide array cloning

CasRx guides were cloned as an array into the pLentiRNAGuide_001 - hU6-RfxCas13d-DR1-BsmBI-EFS-Puro-WPRE plasmid, which was a gift from Neville Sanjana (Addgene plasmid # 138150; http://n2t.net/addgene:138150; RRID:Addgene_138150). To this end, we designed a sequence containing three 23 basepair-long guides interspaced by 2 optimised direct repeats, previously described ^59^ and BsmBI restriction sites to produce compatible overhangs to the entry vector. Such sequence was produced by PCR amplifying a 120 bp-long oligonucleotide (template) using two compatible primers to extend the sequence to its final length (157 bp, including 3 additional nucleotides at each site to favour restriction enzyme activity). Insert and pLentiRNAGuide_001 were linearized with the BsmBI restriction enzyme and ligated using T4 DNA ligase (NEB #M0202S). Primers and templates are provided in Supplementary table 1, part C.

#### PiggyBac plasmids

All piggyBac plasmids used in this study have been obtained with standard molecular cloning techniques using as backbone pSLQ2812 (addgene #84240, a kind gift from the Qi lab). The ecDHFR degron domain was amplified from CAG-DDdCas9VP192-T2A-EGFP-ires-puro (Addgene plasmid # 69534; http://n2t.net/addgene:69534; RRID:Addgene_69534, a gift from Timo Otonkoski). The SMASh degron domain was amplified from pCS6-SMASh-YFP, which was a gift from Michael Lin (Addgene plasmid # 68852; http://n2t.net/addgene:68852; RRID:Addgene_68852). The ER50 degron domain was amplified from pBMN ER50-YFP, a gift from Thomas Wandless (Addgene plasmid # 37259; http://n2t.net/addgene:37259; RRID:Addgene_37259). The AID degron domain was amplified from pcDNA5-H2B-AID-EYFP, a gift from Don Cleveland (Addgene plasmid # 47329; http://n2t.net/addgene:47329; RRID:Addgene_47329). The OsTIR1 protein employed for the AID and mAID degron systems was amplified from pMK232 (CMV-OsTIR1-PURO), which was a gift from Masato Kanemaki (Addgene plasmid # 72834; http://n2t.net/addgene:72834; RRID:Addgene_72834). The KRAB-Split-dCas9 plasmid was generated by inserting the FKBP12^F36V^ domain C-terminally to dCas9 into the plasmid pSLQ2818 pPB: CAG-PYL1-KRAB-IRES-Puro-WPRE-SV40PA PGK-ABI-tagBFP-SpdCas9 (Addgene plasmid # 84241; a gift from Stanley Qi), after exchanging the Puromycin resistance with a resistance for Blasticidin.

The hHDAC4 domain used to generate hHDAC4-dCas9 and dCas9-hHDAC4 repression systems was amplified from pEx1-pEF-H2B-mCherry-T2A-rTetR-HDAC4, which was a gift from Michael Elowitz (Addgene plasmid # 78349; http://n2t.net/addgene:78349; RRID:Addgene_78349). Finally, the CasRx (RfxCas13d) coding sequence was amplified from pLentiRNACRISPR_007 - TetO-NLS-RfxCas13d-NLS-WPRE-EFS-rtTA3-2A-Blast, which was a gift from Neville Sanjana (Addgene plasmid # 138149; http://n2t.net/addgene:138149; RRID:Addgene_138149). Plasmid maps can be found in the Supplementary File 2. Plasmids and their associated sequences are currently being deposited at Addgene.

### RNA extraction, reverse transcription, qPCR

To harvest RNA samples for quantitative PCR (qPCR), ∼ 2 × 10^6^ cells were washed with ice-cold PBS and lysed by directly adding 500 μl of Trizol (Invitrogen). RNA was isolated using the Direct-Zol RNA Miniprep Kit (Zymo Research) following the manufacturer’s instructions. 1μg of RNA were reverse transcribed using Superscript III Reverse Transcriptase (Invitrogen) with random hexamer primers (Thermo Fisher Scientific) and expression levels were quantified in the QuantStudio™ 7 Flex Real-Time PCR machine (Thermo Fisher Scientific) using 2xSybRGreen Master Mix (Applied Biosystems) normalising to Rrm2 and Arpo. All qPCR primers have been validated by PCR and cDNA dilution curve. Primer sequences are listed in Supplementary table 1 part D.

### Live cell flow cytometry

Cell samples for flow cytometry were harvested by washing with PBS, dissociation with trypsin for 7 minutes at 37°C and resuspension in serum-containing medium (DMEM (Sigma), 15% ESC-grade FBS (Gibco), 0.1 mM β-mercaptoethanol). The cell suspension was transferred to a U-bottom 96-well plate, centrifuged for 5 minutes at 500xg at 4°C and resuspended in 70µl flow cytometry buffer (PBS, 10% ESC-grade FBS (Gibco), 0.5mM EDTA) on ice. The samples were analysed using BD FACSCelesta Cell Analyzer (Beckton Dickinson, IC-Nr.: 68186, Serial-Nr.: R66034500035) with 2-Blue6-Violet4-561YG laser configuration, equipped with BD High Throughput Sampler (HTS). The HTS option was used for measurement of fluorescence in 96-well plates. tBFP fluorescence was measured using the 450/440 Band Pass and tRFP or mCherry using the 586/515 Band Pass filters. For the experiments with tRFP and tBFP measurements, violet laser was set to a voltage of 380 and yellow-green laser to 420V. For the experiments involving mCherry and tBFP measurement, the violet laser was set at 340 and yellow-green at 465V. When the HTS option was used, the sample flow-rate was set to 1 or 2 µl/sec and 70% of the total volume in the well was set at sample volume. At least 10,000 events were recorded. When the cells were analysed using FACS tubes, at least 30,000 events were recorded.

## Data analysis

### Flow cytometry data analysis

Data analysis of flow cytometry files was performed using the R programming language and the packages “FlowCore”, “OpenCyto” and “ggcyto”. In order to clearly distinguish events corresponding to single, live cells, two gates are applied sequentially. First, events were gated based on the Forward and Side Scatter Area (FSC-A and SSC-A) using openCyto:::.boundary() in order to select events corresponding to live cells. Second, to separate single cells (singlets) from doublets, the events showing a linear relationship between Forward Scatter Height and Area (FSC-H and FSC-A) were automatically selected using openCyto:::.singletGate(). Importantly, the same gating coordinates were applied to all files within one experiment and, where possible, across experiments. The experiments involved either tBFP and tRFP as fluorochromes or tBFP and mCherry. Because the overlap between emission spectra is minimal, no compensation was deemed to be necessary.

The fluorescence distributions were plotted by extracting the relevant parameters of the singlet-gated populations using “ggplot2” and the ggridges package with the function geom_density_ridges(), which calculates density estimates from the provided data. To quantify protein abundance from flow cytometry data, the Median Fluorescence Intensity (MFI) was calculated for a given sample, followed by correction for the cells’ autofluorescence through subtraction of the MFI of one or multiple non-fluorescent control samples of the same cell line.

#### Analysis of degron properties

To analyse the properties associated with the different degrons, each sample was corrected for autofluorescence background by taking the mean of the MFI of non-fluorescent cells treated with the maximal concentration of ligand and mock-treated. We then calculated the tRFP/tBFP ratio and normalised it to the mean of the tRFP/tBFP ratio for the no-degron control cell line, averaged across all replicates treated with maximal ligand concentration and mock-treated samples. This relative tRFP/tBFP ratio was then expressed as a percentage. To quantify degron properties, we then defined the tRFP/tBFP ratio at the stabilised state as A and the ratio under destabilised conditions as B.

The “degradation leakiness”, which measures background destabilisation of the stabilised state (e.g. destabilisation conferred by the degron tag *per se* in the absence of degrader), was defined as:

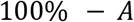

The “degradation efficiency”, which measures the reduction of protein abundance relative to the no-degron control, was defined as:

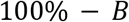

The “dynamic range” for each degron-dCas9 construct, which measures the range of protein concentrations that can be achieved by modifying the ligand concentration, is calculated as *B*/*A*.

### Modelling the dynamics of repression and reversibility

Curve fitting and deterministic modelling were performed using the R programming language and the packages “minpack.lm” for non-linear least square (NLS) problems computation and “deSolve” for solving ordinary differential equations.

We described the system with two ordinary differential equations (ODEs), where equation (1) describes the expression of the Cas-Repressor R and equation (2) the target gene expression T, the production of which is modulated by the Cas-Repressor:

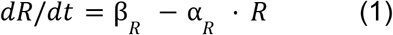

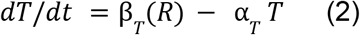

Where α and β denote the degradation and production rates, respectively. β_R_ and α_*T*_ are assumed to be constant, while α_*R*_ is modulated by the dTAG-13 concentration and the effective production rate of the target β_T_(R) is a function of the repressor levels.

#### Quantification of repressor half times

Since α_*R*_ is modulated by the dTAG-13 concentration, we estimated the parameter both in the presence (α_*R*+_) and in the absence (α_*R*−_) of the ligand from time course data upon dTAG-13 addition and withdrawal, respectively. The background-corrected MFI values were rescaled between 0 and 1. For the dTAG withdrawal time course, where all repressors reached their maximal level in less than 25h, the mean MFI of the 25h to 150h time points was set to 1 and the mean MFI at time 0h to 0. To calculate the time required for the Cas-Repressor to reach half of its maximal concentration, the time course was fit with the analytical solution of eq. (1) with the initial condition R(0) = 0:

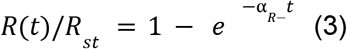

Where *R*_*st*_ denotes the level of R at steady state (t>24h) and *R*(*t*)/*R*_*st*_ represents the scaled Cas-Repressors expression level. Since the half-time required for the Cas-Repressor to reach the steady state is given by 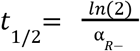, we substituted α_*R*−_ in eq. (3) with *In*(2)/*t*_1/2_ to directly compute the half-time and the associated standard error for each Cas-Repressor. To estimate the repressor degradation rate and associated half time in the presence of dTAG-13, the Cas-Repressor downregulation time course upon dTAG-13 addition was rescaled such that the mean MFI of the 25h to 150h time-points, where all repressor constructs had been fully degraded, was set to zero and the mean MFI before the treatment was set to 1. The data was then fit with the analytical solution of eq. (1) with the initial condition *R*_0_ =1 and a steady state of 0:

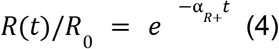

Where α_*R*+_ is the degradation rate of the Cas-Repressor upon dTAG-13 addition and the half time of repressor removal was estimated as described above.

#### Parameter estimation of repressor-dependent target repression

In order to quantify the relationship between Cas-Repressor concentration and target repression β_*T*_ = *f*(*R*), we used the dTAG-13 titration data sets with 4 days of treatment, assuming that the systems had reached a steady state at that time. Background-corrected MFI values of Cas-Repressors (tBFP) were scaled between 0 and 1 (min-max scaling), while the target gene (mCherry) was computed as a fold change relative to levels in the absence of repression (500nM dTAG-13). Describing the dose-response relationship between the repressor and the target with a Hill function, we then solved eq. (2) at steady state:

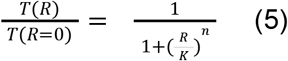

Where 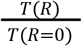 describes the scaled target levels, K denotes the levels of R that leads to 50% target repression and n represents the Hill coefficient. K and n were estimated for each repressor construct.

#### Estimation of the target degradation rate

To estimate the degradation rate of the target gene α_*T*_, we used the mCherry time course upon ligand addition in the CasRx cell line. Since the repressor was fully degraded in <1h and target repression can be assumed to be immediate in this system, which relies on mRNA degradation, the dynamics of reporter upregulation can be assumed to be uniquely determined by the mCherry degradation rate. After normalising the background-subtracted MFI of the target gene to the mean MFI at time-point 150h, we estimated the mCherry degradation rate by fitting the analytical solution of eq. (2) in the absence of repression to the data:

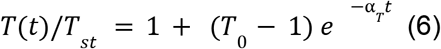

Where *T*_*st*_ is the final target level (at 150h) and *T*_0_ the initial target level. The mCherry degradation rate α_*T*_, was estimated to be 0.036h^-1^.

#### Quantification of repression delay

To understand whether target repression and derepression is immediate or follows changes in repressor levels with a delay, we used the ODE model to simulate target expression with the estimated parameters, either assuming an immediate (Δ*t*=0) or a delayed (Δ*t*>0) effect of the repressor on the target. Different values were tested for Δ*t*, ranging from 0 to 25h and numerical solutions were compared to the experimentally measured mCherry levels through computing the Mean Absolute Error (MAE). The Δ*t* that minimised the MAE is reported.

### Modelling of NANOG - target genes dose-response curves

To quantify how known NANOG target genes *Sox2, Klf4, Rex1, Nrb01* and *Xist* responded to NANOG perturbations, we fitted a Hill function to the dose-response curves. To this end, target gene expression was first normalised to the NANOG-high state (500nM dTAG-13). Then we used the “minpack.lm” package in R, to fit the dose-response curve with a 4 parameter Hill-type function using a non-linear least square approach using the formula:

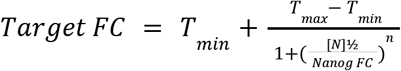

The equation describes the fold change variation of the target (*Target FC*) as a function of *Nanog* fold change (*Nanog FC*). *T*_*max*_ is the target fold change at 500nM dTAG-13 and is set equal to 1, while *T*_*min*_ is the target fold change when *Nanog* is 0 and is estimated by fitting the data with the NLS approach. Through this, we can extract [*N*]½, which corresponds to the *Nanog* fold change that leads to half of the maximal target variation, and the Hill coefficient *n*, which indicates the degree of non-linearity.

For genes repressed by NANOG, namely Xist in this case, the equation is written as:

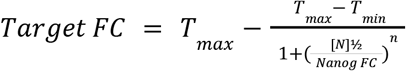

with *T*_*min*_ set equal to 1.

## Acknowledgments

We thank Oriana Genolet for generating the 1.8XX Esrrb-mCherry and Nanog-mCherry cell lines. We also thank Liat Ravid Lustig for scientific advice and Francesca Rossi for scientific discussion. We thank the Max Planck Institute for Molecular Genetics FACS facility. This work was supported by the Max-Planck Research Group Leader program, E:bio Module III—Xnet grant (BMBF 031L0072) and Human Frontiers Science Program (CDA-00064/2018) to E.G.S. G.N. was supported by the European Union’s Horizon 2020 Research and Innovation Program (Marie Skłodowska-Curie ITN PEP-NET).

## Author contributions

GN and EGS conceived the project and designed the experiments. RAFG cloned most of the degron-dCas9-tRFP-P2A-tBFP and no-degron control plasmids. GN cloned all other plasmids, performed the experiments and analysed the data. GN and EGS wrote the manuscript.

## Supplementary Figures

**Fig. S1:**
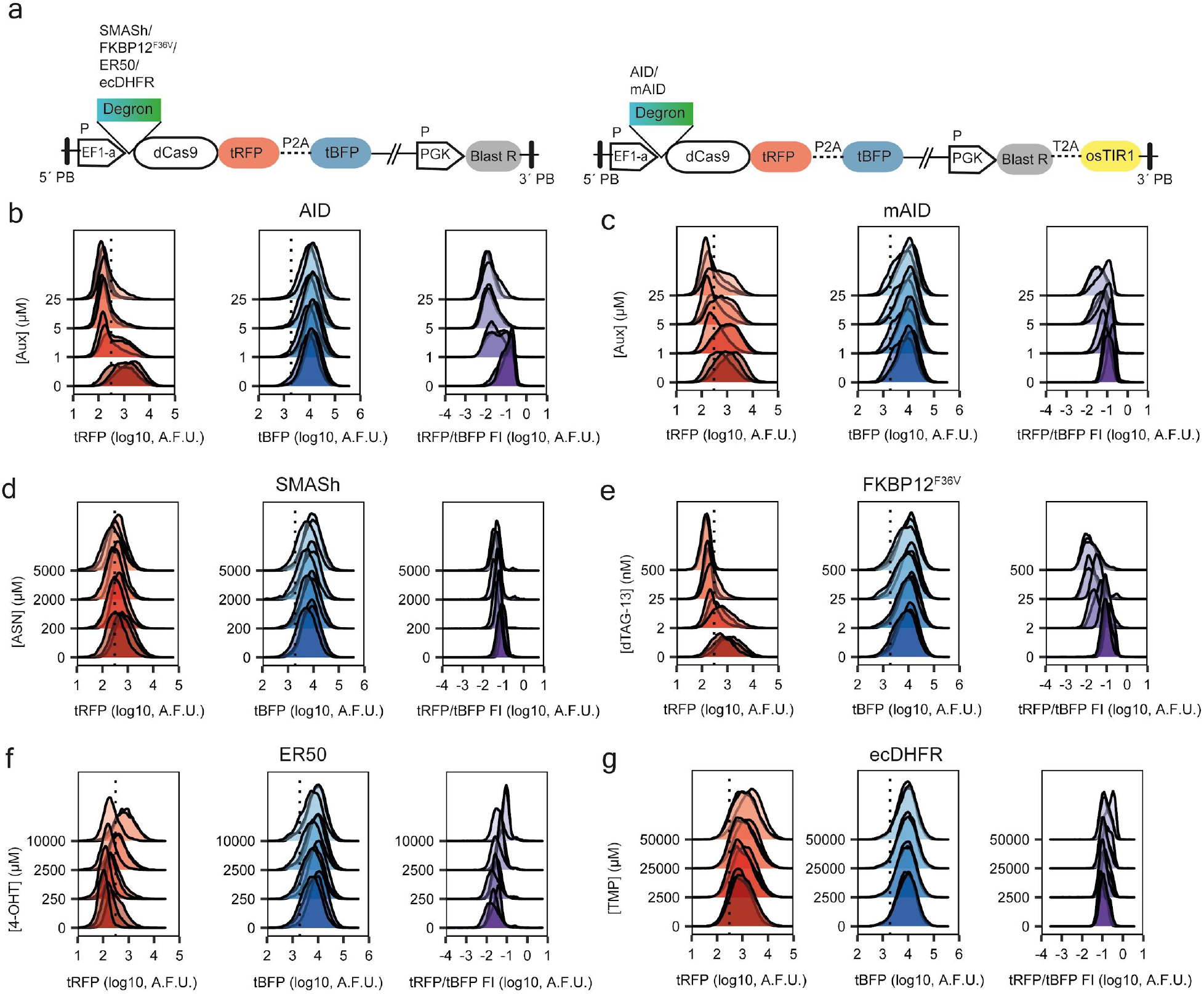
Comparison of degron domains to control dCas9. **a**, A detailed schematic of degron-dCas9-tRFP-P2A-tBFP plasmids. The region between the 5’ and 3’ PB sites is genomically integrated. The degrons SMASh, FKBP12^F36V^, ER50 and ecDHFR (left) do not require any accessory protein for their degradation. The blasticidin resistance gene is expressed under a separate promoter (PGK). The no-degron control plasmid is based on the same design. The AID and mAID degrons (right) require the TIR1 protein from Oryza sativa (OsTIR1), which is expressed under the same promoter of the blasticidin resistance, separated by a T2A site. **b-g**, Density plots showing the fluorescence distributions across cells for tRFP, tBFP and the tRFP/tBFP ratio at different ligand concentrations for each degron domain as indicated. Three biological replicates for each concentration are overlaid. A.F.U. Arbitrary Fluorescence Units.

**Fig. S2:**
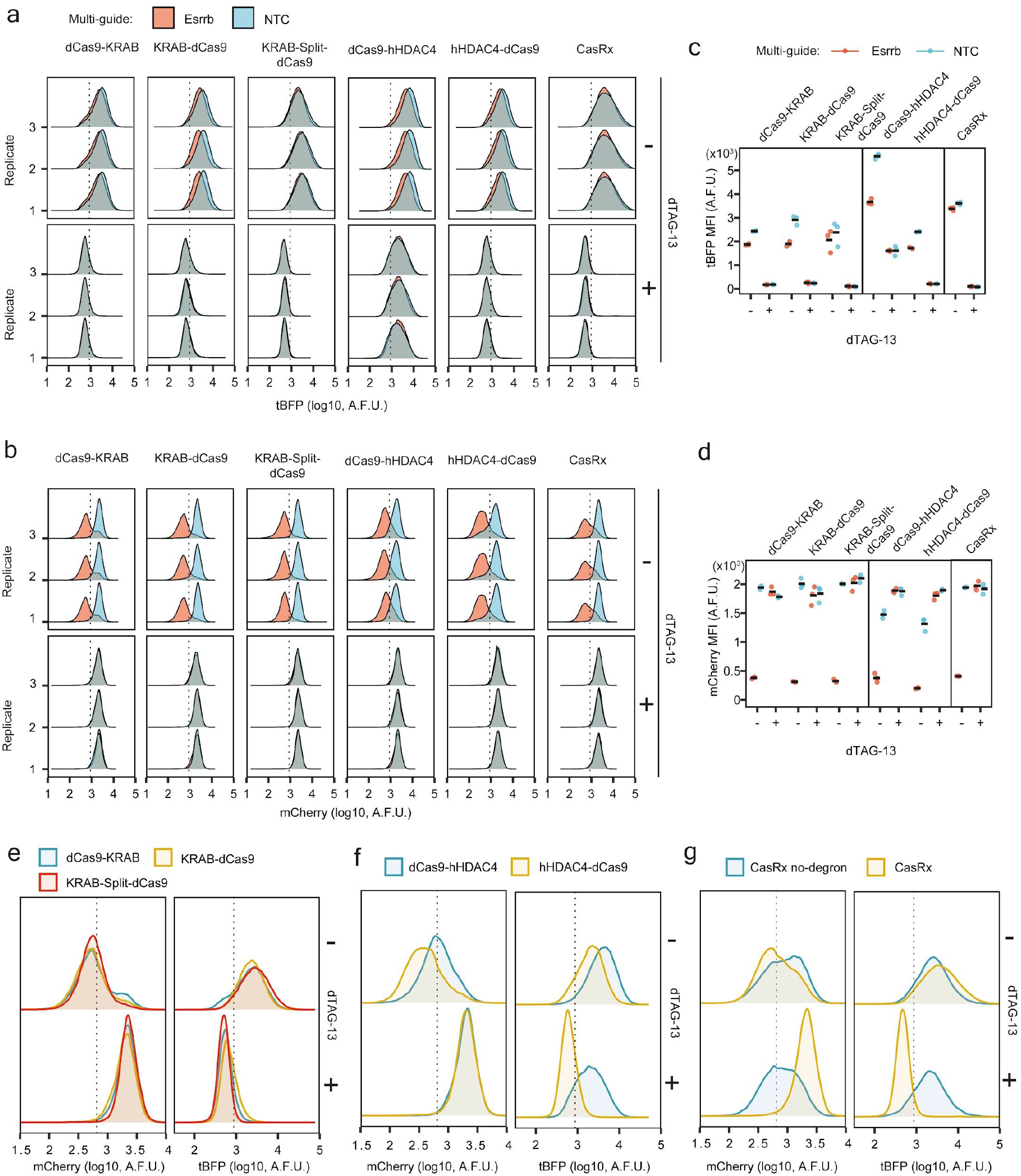
Testing inducibility and efficiency of repression through degron-Cas-repressors in an endogenous reporter system. **a**,**b**, Density plots of tBFP (**a**) or mCherry levels (**b**) in cells containing Esrrb- (red) or non- (blue) targeting guides, in -dTAG-13 (top) or +dTAG-13 (bottom) conditions, measured by flow cytometry. **c**,**d**, Background-subtracted MFI of tBFP (**c**) or mCherry (**d**) for degron-Cas-repressor cell lines indicated on top, in - or +dTAG-13 conditions. For Esrrb- (red) or non- (blue) targeting guides. Dots showing 3 biological replicates and horizontal bars their mean. **e**-**g**, Density plots showing the overlaid distributions for mCherry (left) and tBFP (right), in -dTAG-13 (top) and +dTAG-13 condition (bottom) in cells expressing Esrrb-targeting guides for the three KRAB-mediated repression systems (**e**) the two hHDAC4-based systems (**f**) and the CasRx system (**g**). For the degron-CasRx construct, the same construct lacking the degron domain (CasRx no-degron) was included for comparison. The three biological replicates are merged together. In **a**,**b**,**e**-**g** the dotted line indicates the 99th percentile of non-fluorescent control cell line. A.F.U.= arbitrary fluorescence units.

**Fig. S3:**
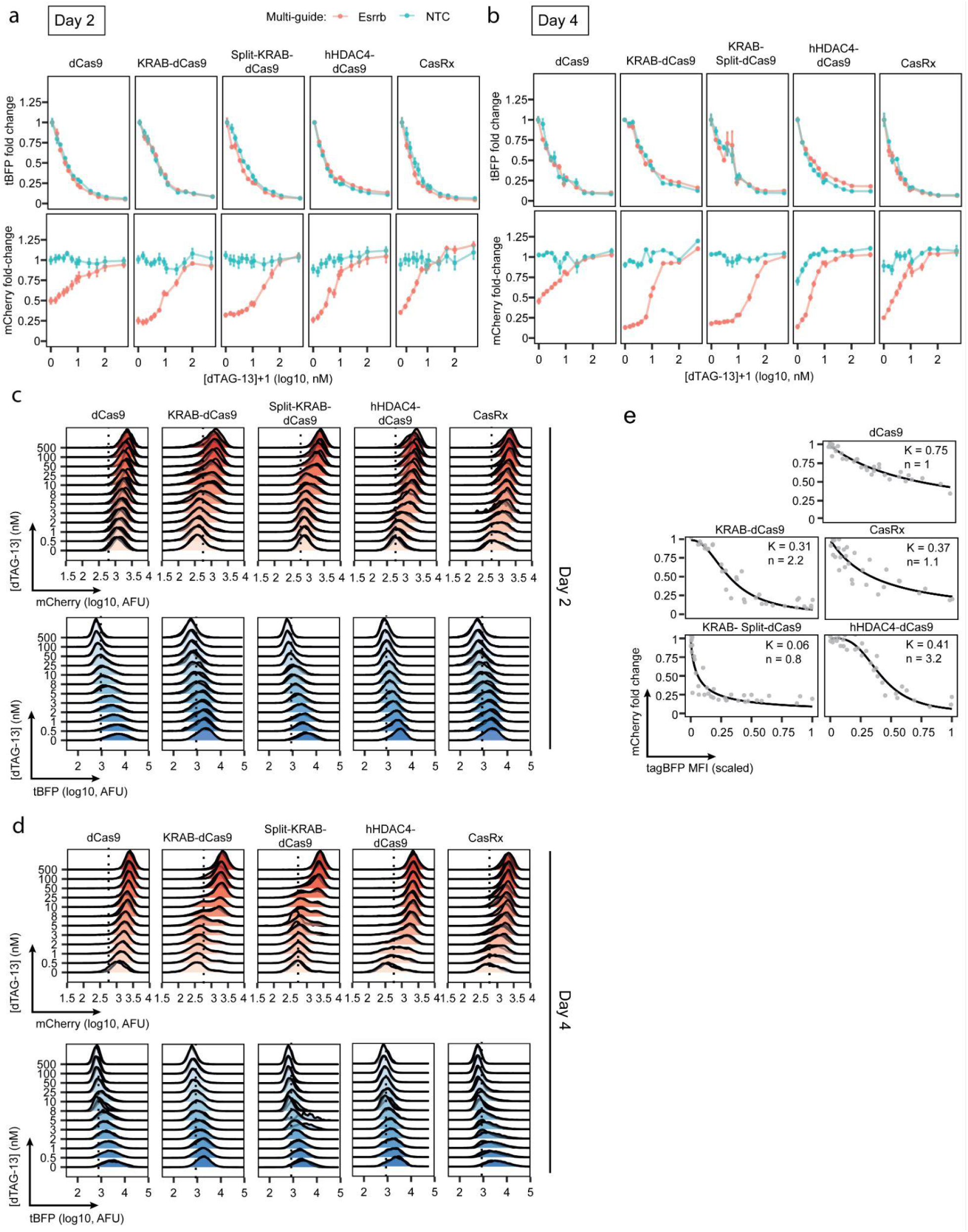
Assessing tunability of degron-Cas-repressor systems. **a, b**, tBFP (top) and mCherry (bottom) fold change of background subtracted MFI in cells expressing Esrrb-targeting guides (red) and cells expressing NTC guides, at different dTAG-13 concentrations (y-axis, log10 +1 scaled, in nM) after 2 (**a**) or 4 (**b**) days of treatment. The fold change is calculated relative to the mean of the MFI in cells with non-targeting guides at the different dTAG-13 concentrations. The mean of three biological replicates (dots) ± standard deviation (vertical bars) is shown. Lines connect the means. **d, c**, Density plots of mCherry (top) or tBFP (bottom) expression levels measured by flow cytometry at increasing dTAG-13 concentrations (y-axis, in nM) for degron-Cas-repressors indicated on top in cells expressing Esrrb-targeting guides at 2 (**c**) and 4 (**d**) days of treatment. Three biological replicates are overlaid. dCas9 and KRAB-Split-dCas9 are the same construct (KRAB-Split-dCas9) without or with addition of 100µM ABA, respectively. **e**, Hill function fitted to titration experiments at the 4 day time point. Shown is the mCherry fold change (y-axis) at varying tBFP levels (min.-max. scaled, x-axis) for all replicates and dTAG-13 concentrations (grey dots) and the best fitting Hill-type equation (black line) resulting from a non-linear least square (NLS) approach to find the parameters K (repression coefficient) and n (Hill coefficient), which are reported inside each plot.

**Fig. S4:**
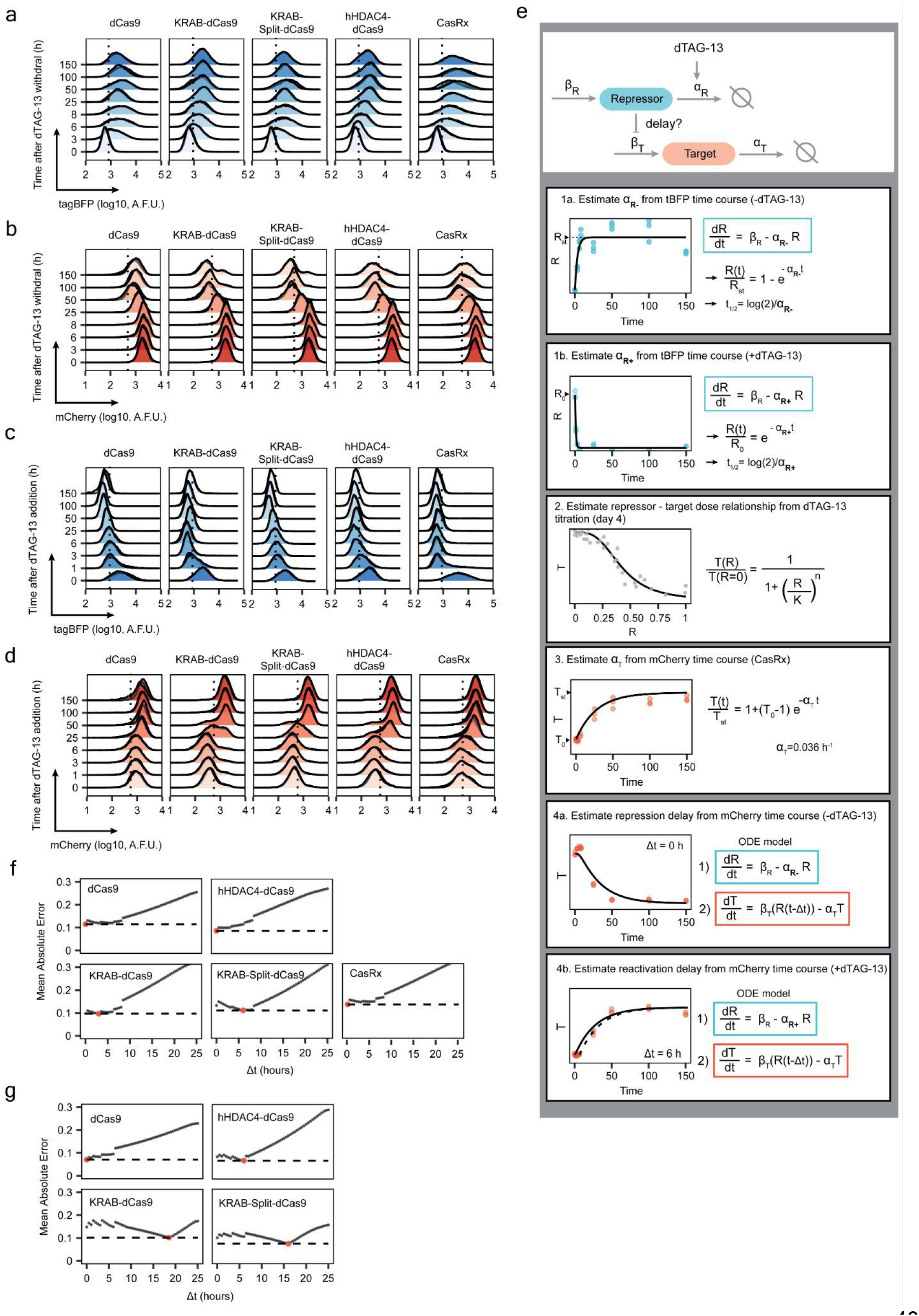
Measuring the dynamics of repression and reversibility associated with degron-Cas-repressor systems. **a, b**, tBFP (**a**) and mCherry (**b**) levels measured by flow cytometry (x axis) at different time-points (y axis, in hours from dTAG-13 withdrawal) for the experiment performed to measure the dynamics of repression. Different repressor systems are indicated on top. Three biological replicates are overlaid. **c, d** tBFP (**c**) and mCherry (**d**) levels measured by flow cytometry (x axis) at different time-points (y axis, in hours from dTAG13 addition) for the experiment performed to measure the dynamics of repression reversibility. Different repressor systems are indicated on top. Three biological replicates are overlaid. **e**, Modelling approach used to quantify the dynamic of target gene repression and derepression for the different degron-Cas-repressors. Repressor (R) and Target gene (T) dynamics were simulated with an ODE model, describing production with rate and degradation with rate α for both R and T. In step 1, repressor degradation rates in the presence (α_R+_) and absence (α_R-_) of dTAG-13 and the resulting half life (t_1/2_) were estimated by fitting the tBFP time course in the presence (1b) or absence (1a) of dTAG-13 with the equation describing the repressor as indicated. In step 2, the relationship between target gene expression and repressor level at the steady state is modelled by fitting an Hill curve to the titration data from Fig. S3e. In step 3, the mCherry derepression time course of the CasRx cell line is used to estimate the mCherry degradation rate (α_T_). All the parameters estimated in steps 1-3 are then combined in step 4a and 4b to simulate a model of two ODEs to describe the dynamic of repression and derepression, respectively. By comparing experimental data with simulations, assuming different delays (Δt) between repressor up- or downregulation and effects on target gene expression, the value for Δt that could best reproduce the data was identified. **f**, Mean absolute error (MAE) for the simulations of the ODE model used for studying the repression dynamic of different degron-Cas-repressors, for different Δt. Each grey dot represents the MAE of the simulation for the corresponding Δt. The Δt associated with the lowest MAE is marked with a red dot and the dashed line indicates the corresponding MAE. **g**, As in (**f**), but for the models used to simulate the reversibility dynamics. Here the MAE is not shown for CasRx because the CasRx experiment was used to calculate the degradation rate of the target protein ESRRB-mCherry.

**Fig. S5:**
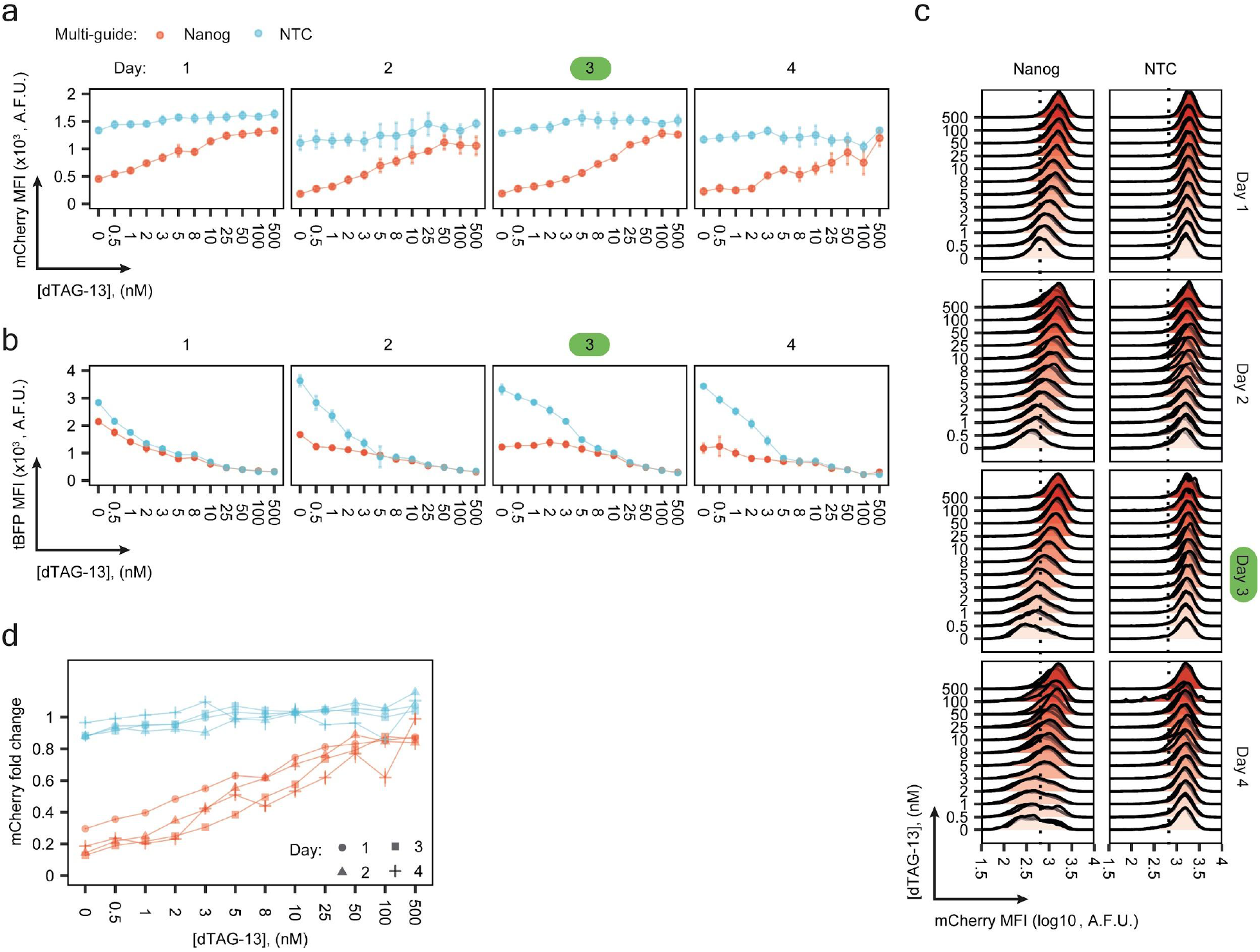
NANOG dose-response curves measured using CasTuner. **a, b**, Background subtracted MFI for mCherry (**a**) or tBFP (**b**) in Nanog-mCherry mESCs expressing the hHDAC4-dCas9 CasTuner system and Nanog- (red) or non- (blue) targeting guides, at different dTAG-13 concentrations (indicated on x-axis). The 4 panels from left to right represent measurement performed after 1 to 4 days from induction of knock-down. Dots show the mean of three biological replicates and vertical lines the standard deviation. Also lines connecting the mean values are shown. **c**, mCherry levels measured by flow cytometry in cells treated with the indicated dTAG-13 concentrations (y-axis) with Nanog-targeting guides (left) or NTC guides (right). Data from measurements at day 1 to 4 from knock-down induction, as indicated on the right. In (**a**-**c**) the time-point used for analysing Nanog dose-response curves is marked in green. **d**, Nanog-mCherry fold change in cells expressing targeting (red) or non-targeting guides (blue), at different dTAG-13 concentrations (x axis) measured by flow cytometry. Symbols indicate different measurement time points as indicated. AFU=arbitrary fluorescence units.

## Supplementary Files

**Supplementary Table 1**. Reagents

Part A: Sequences of sgRNA for dCas9.

Part B: Sequences of sgRNA for CasRx.

Part C: Oligonucleotides sequences for cloning of CasRx guides.

Part D: qPCR primers.

**Supplementary File 1**. Strategy for double-sorting by FACS of cell lines with degron-Cas-repressor systems.

**Supplementary File 2**. Maps of plasmids used in this study.

