## Supplementary File 1 for "CasTuner: a degron and CRISPR/Cas-based toolkit for analog tuning of endogenous gene expression"

Description: Strategy for double sorting of degron-Cas-Repressor systems
Cells are first sorted gating on BFP levels with a wider gate (BFP+), expanded and then sorted again (M population).
*Page 2:  Example of first sorting. Sample: degron-dCas9-KRAB. Population sorted: BFP+
*Page 3: Example of second sorting. Sample: degron-dCas9-KRAB. Population sorted: M and Hi. Population M is used for all the experiments described in the manuscript.
The same gating coordinates are applied to all samples.

### BD FACSDiva 8.0.1

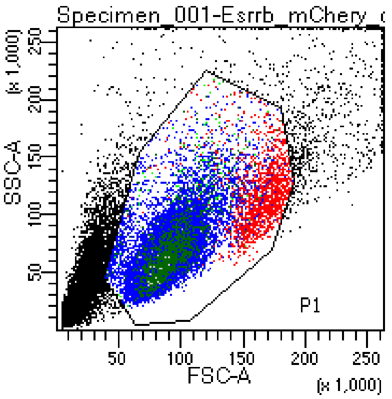


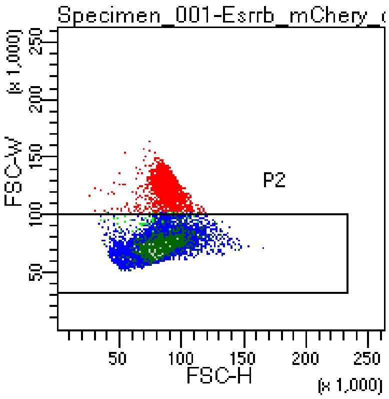


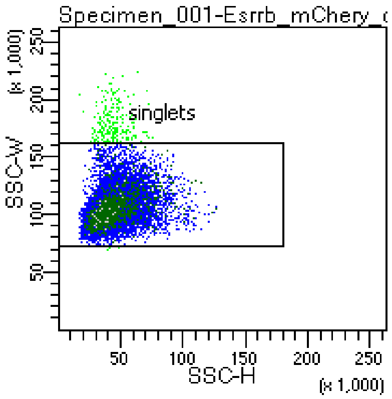


Tube: Esrrb_mChery_dCas9_dTAG_KRAB_BFP

| Population | #Events | %Parent | %Total |
|---|---|---|---|
| All Events | 37,981 | #### | 100.0 |
| P1 | 30,000 | 79.0 | 79.0 |
| P2 | 28,317 | 94.4 | 74.6 |
| singlets | 28,078 | 99.2 | 73.9 |
| BFP+ | 1,369 | 4.9 | 3.6 |
| negative | 27 | 0.1 | 0.1 |

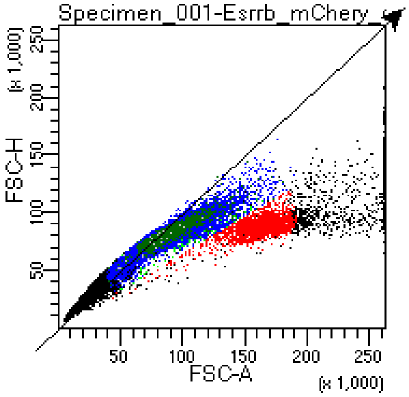


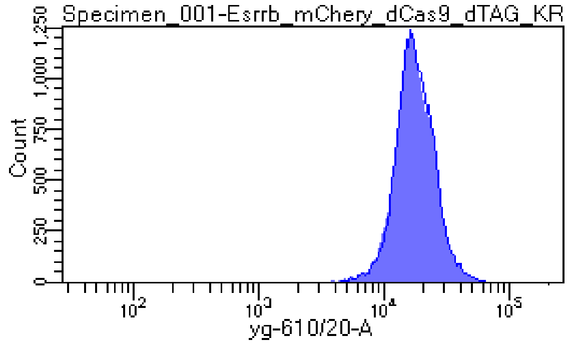


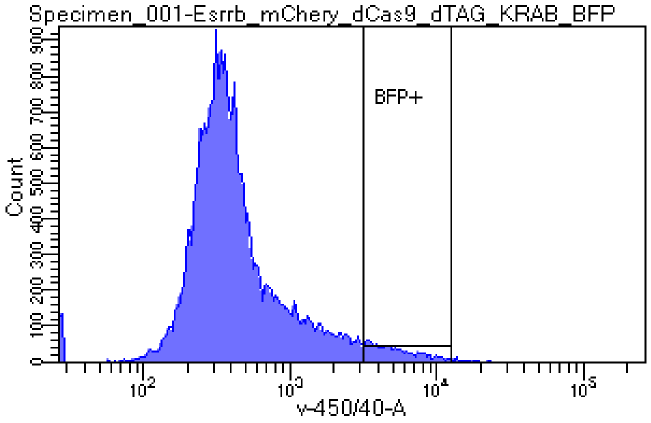

### BD FACSDiva 8.0.1

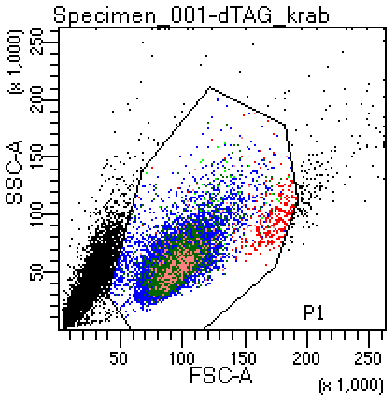


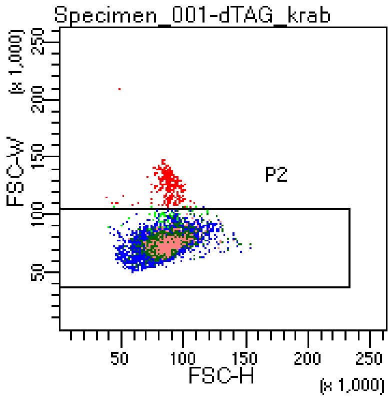


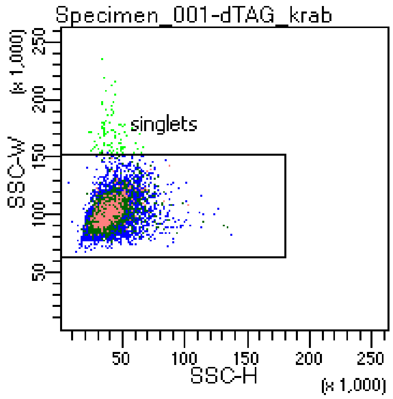


Tube: dTAG_krab

| Population | #Events | %Parent | %Total |
|---|---|---|---|
| All Events | 24,131 | #### | 100.0 |
| P1 | 20,000 | 82.9 | 82.9 |
| P2 | 19,683 | 98.4 | 81.6 |
| singlets | 19,594 | 99.5 | 81.2 |
| M | 2,645 | 13.5 | 11.0 |
| Hi | 363 | 1.9 | 1.5 |

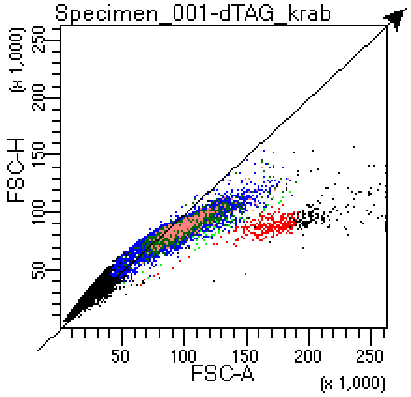


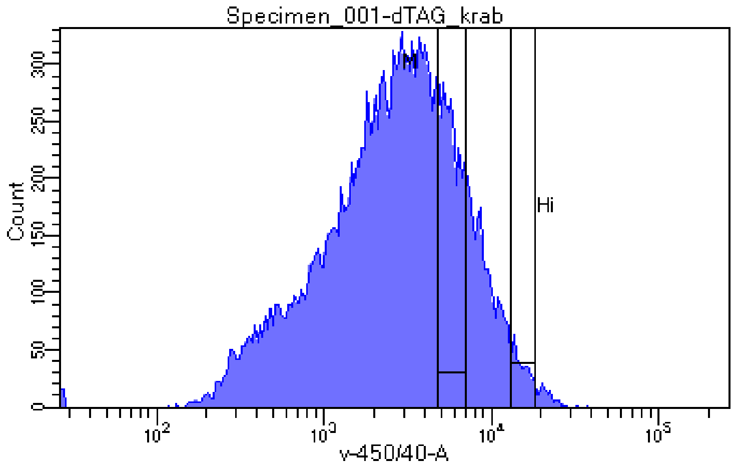


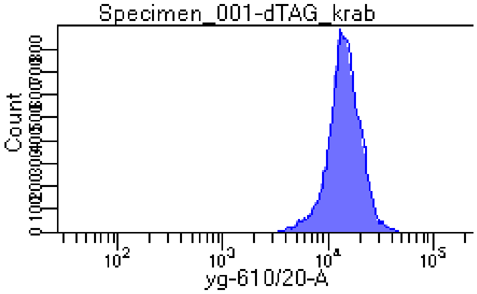
