## Supplementary File 2 for "CasTuner: a degron and CRISPR/Cas-based toolkit for analog tuning of endogenous gene expression"

Short name:
dCas9-tagRFPt-P2A-tagBFP
Full name:
pSLPB2-SpdCas9-tagRFPt-P2A-tagBFP-PGK-Blasticidin

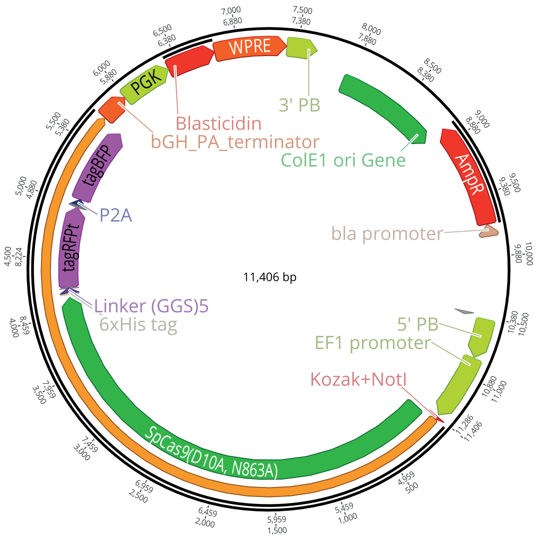


Short name:
AID-dCas9-tagRFPt-P2A-tagBFP
Full name:
pSLPB3B-AID-SpdCas9-tagRFPt-P2A-tagBFP-PGK-Blasticidin-T2A-osTIR1

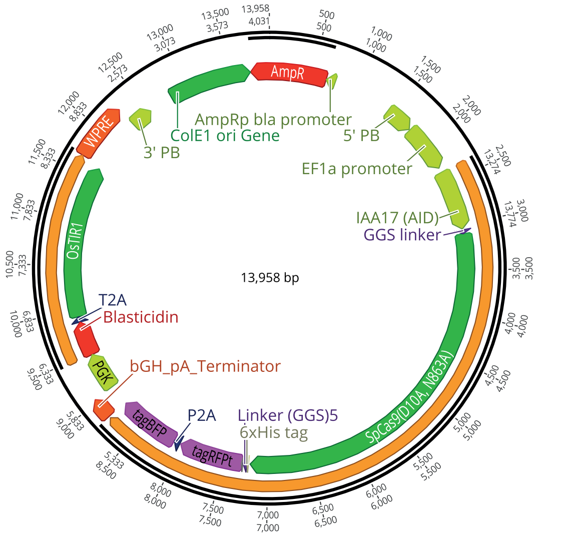


Short name:
mAID-dCas9-tagRFPt-P2A-tagBFP
Full name:
pSLPB3B-AID-SpdCas9-tagRFPt-P2A-tagBFP-PGK-Blasticidin-T2A-osTIR1

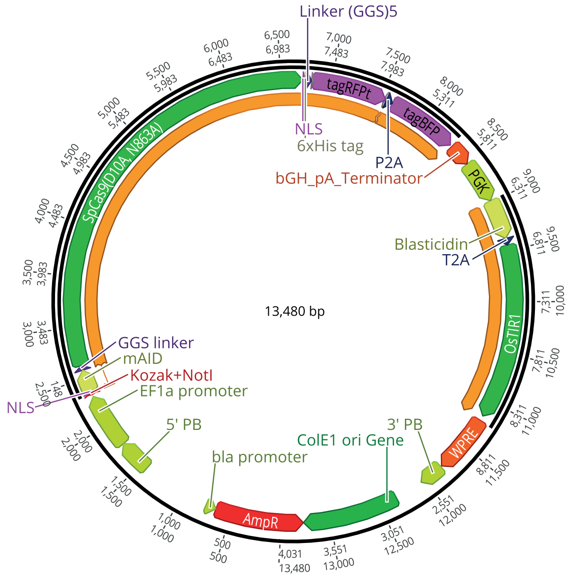


Short name:
ER50-dCas9-tagRFPt-P2A-tagBFP
Full name:
pSLPB2B-ER50-SpdCas9-tagRFPt-P2A-tagBFP-PGK-Blasticidin

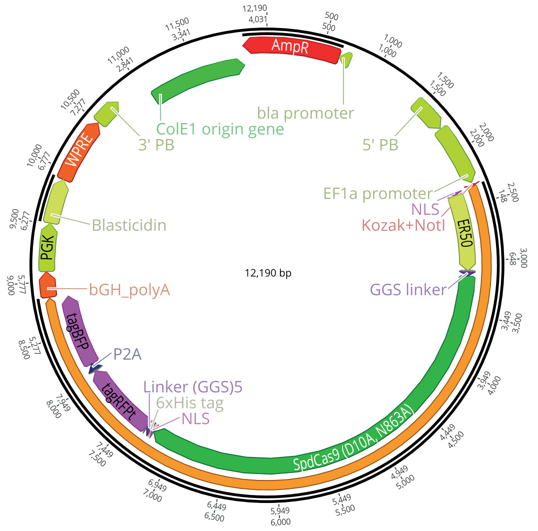

Short name:
SMASh-dCas9-tagRFPt-P2A-tagBFP
Full name:
pSLPB2B-SMASh-SpdCas9-tagRFPt-P2A-tagBFP-PGK-Blasticidin

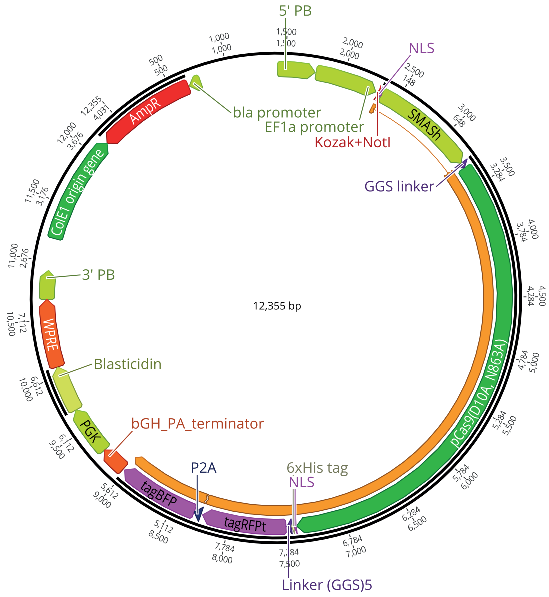


Short name:
ecDHFR-dCas9-tagRFPt-P2A-tagBFP
Full name:
pSLPB2B-ecDHFR-SpdCas9-tagRFPt-P2A-tagBFP-PGK-Blasticidin

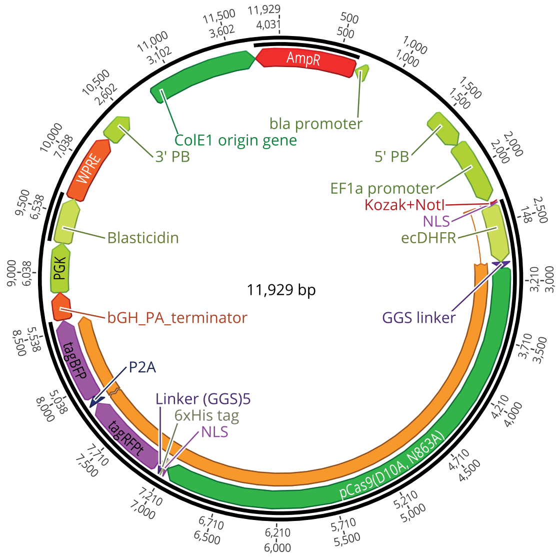


Short name:
FKBP12_F36V-dCas9-tagRFPt-P2A-tagBFP
Full name:
pSLPB2B-FKBP12_F36V-SpdCas9-tagRFPt-P2A-tagBFP-PGK-Blasticidin

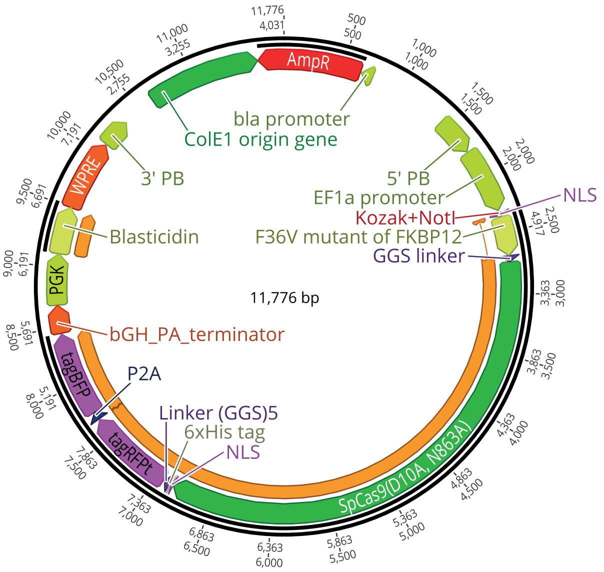


Short name:
FKBP12_F36V-CasRx-tagBFP
Full name:
pSLPB2B-FKBP12_F36V-CasRx-tagBFP-PGK-Blasticidin

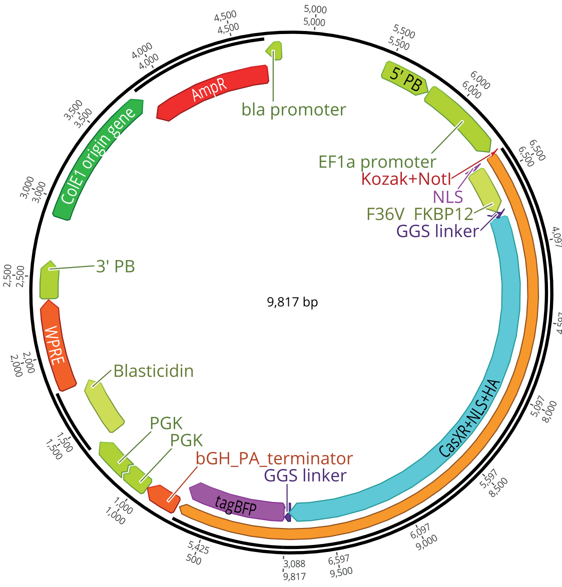

Short name:
FKBP12_F36V-dCas9-KRAB-tagBFP
Full name:
pSLPB2B-FKBP12_F36V-SpdCas9-KRAB-tagBFP-PGK-Blasticidin

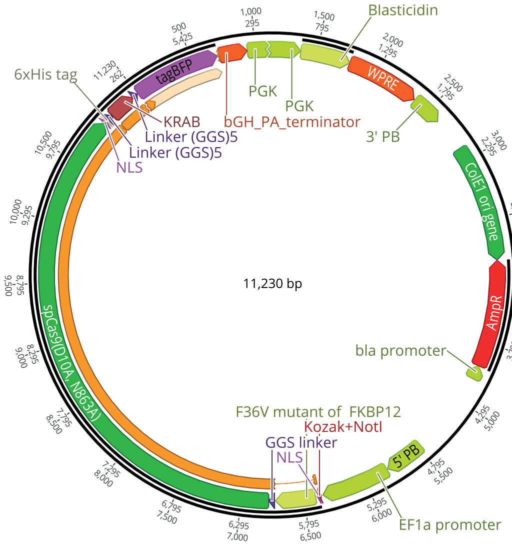


Short name:
FKBP12_F36V-dCas9-hHDAC4-tagBFP
Full name:
pSLPB2B-FKBP12_F36V-SpdCas9-hHDAC4-tagBFP-PGK-Blasticidin

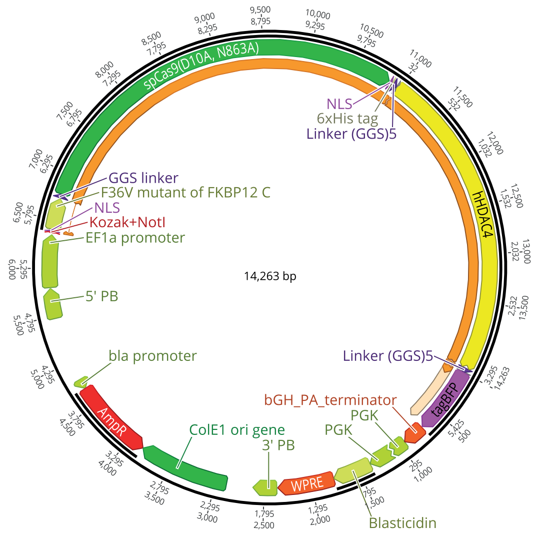


Short name:
FKBP12_F36V-KRAB-dCas9-tagBFP
Full name:
pSLPB2B-FKBP12_F36V-KRAB-SpdCas9-tagBFP-PGK-Blasticidin

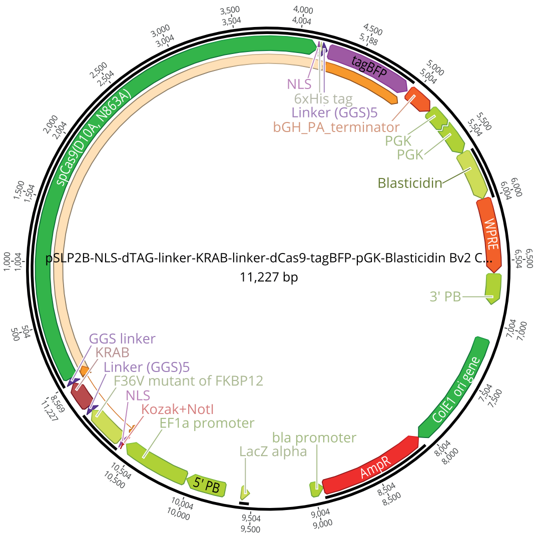


Short name:
FKBP12_F36V-hHDAC4-dCas9-tagBFP
Full name:
pSLPB2B-FKBP12_F36V-hHDAC4-SpdCas9-tagBFP-PGK-Blasticidin

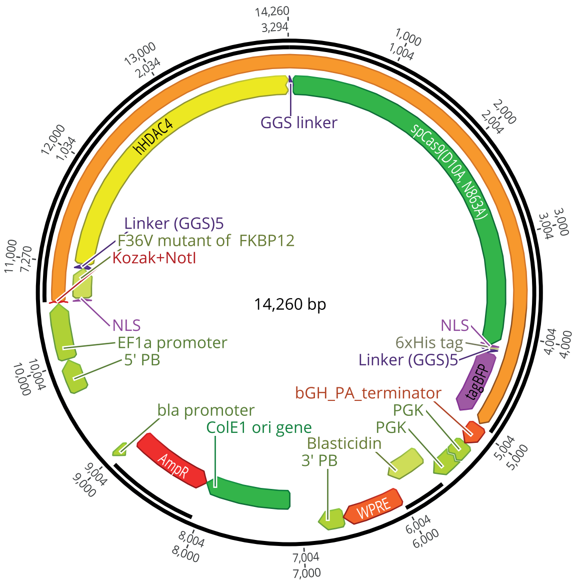

Short name:
KRAB-Split-dCas9-FKBP12_F36V
Full name:
pSLQ2818_Blasti_pPB: CAG-PYL1-KRAB-IRES-Blasti-WPRE-SV40PA
PGK-ABI-tagBFP-SpdCas9-FKBP_F36V

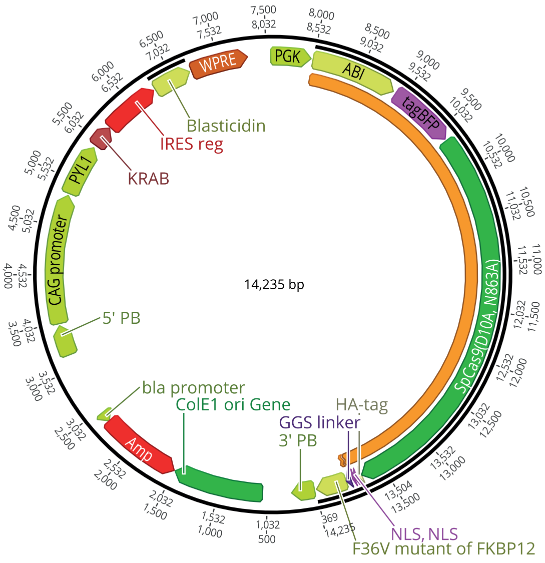
